# Autoencoder-based anomaly mapping and visualization of white matter deformation with application to glioma cases

**DOI:** 10.64898/2026.10.08.757575

**Authors:** Boshra Shams, Lucius S. Fekonja, Peter Vajkoczy, Thomas Picht, Dogu Baran Aydogan

## Abstract

**Introduction:** Gliomas can alter white matter pathways, causing structural deformation and functional impairments. However, there is no method to map these structural alterations across the whole brain to support diagnosis, prognosis, and treatment planning. To address this gap, we propose a novel framework based on a convolutional autoencoder that learns geometry-preserving streamline embeddings for mapping structural connectivity anomalies

**Methods:** We trained and validated the autoencoder on 100 million synthetic streamlines and evaluated the anomaly-mapping pipeline using data from 118 healthy subjects. We analyzed how different tractography algorithms affect anomaly scores and demonstrated its feasibility in five glioma patients using multi-shell and single-shell diffusion MRI.

**Results:** Latent-space distances strongly associated with native-space geometric distances (*R*² = 0.82–0.98), supporting preserved streamline geometry in the learned embeddings. Anomaly scores varied with both the input and reference tractography methods. Patient anomaly maps showed elevated scores near gliomas and extending distally along affected pathways. These whole-brain anomaly maps were generated within minutes.

**Conclusion:** The framework offers a practical way to quantify and visualize geometric deviations in whole-brain tractograms, providing novel insights beyond current network and microstructural analyses, with relevance to clinical applications where white matter deformation is pertinent.

## 1. Introduction

Gliomas are highly infiltrative tumors that invade surrounding tissue (Louis et al., 2021) and can affect the brain’s white matter at both micro- and macrostructural levels, through infiltration and mass effect (Ferrer et al., 2018; Lipková et al., 2022). These processes can compromise fiber tract integrity, leading to tissue degeneration and deformation (Brooks et al., 2021). In addition, tumor mass exerts mechanical forces—solid stress and tumor-associated edema—which can deform and compress brain structures (Nia et al., 2016). These can contribute to neurological and cognitive deficits (Fekonja et al., 2021, 2022; Prasse et al., 2023; Seano et al., 2019; Shams et al., 2022, 2023). Functional imaging studies have demonstrated that gliomas can alter brain function not only in the vicinity of the lesion but also in distant regions, including the contralateral hemisphere (Duffau, 2014, 2020; Stoecklein et al., 2020). Therefore, studying and mapping white matter deformation is crucial for understanding a glioma’s impact on the brain’s structure and function. Developing reliable and robust methods to characterize underlying white matter structural alterations is essential for establishing stronger links between structural abnormalities and functional impairments. To the best of our knowledge, however, there is no established method for this purpose. Valuable white matter deformation information can benefit clinical assessments and aid in preoperative planning for surgical interventions.

The brain’s white matter, which primarily contains axonal connections, can be studied using diffusion MRI (dMRI)-based tractography. This technique uniquely enables the non-invasive, in vivo mapping of human brain connections (Jeurissen et al., 2019). Tractography has become essential for studying white matter structure in healthy and diseased brains, offering insights into how conditions like gliomas impact structural connectivity (D’Souza et al., 2021; Fekonja et al., 2022; Shams et al., 2022, 2023; Silvestri et al., 2022; Wei et al., 2023). It informs about the three-dimensional (3D) geometry of connections within the white matter. Analyzing the individual connections, i.e., streamlines, offers an opportunity to study deformations. By quantifying deviations from the typical connectivity patterns observed in healthy individuals, tractography enables characterization of structural deformations. This information can be used to rank the severity and spatial extent of white matter deformation, providing previously unavailable insights into the impact of pathologies on the brain.

A systematic study of deformations requires a comprehensive set of reference streamlines that represent typical, healthy white matter connections. However, the inherent complexity and variability of brain connectivity demand large whole-brain tractograms, often containing millions of streamlines per individual (Fekonja et al., 2022; Reid et al., 2020; R. E. Smith et al., 2022). This scale presents significant computational challenges, as processing such datasets, including tasks like streamline filtering, bundle labeling, and analyzing population studies, can be resource-intensive and time-consuming (Chandio et al., 2020; Garyfallidis et al., 2018; Gupta et al., 2018; Legarreta et al., 2021). To address these challenges, deep learning has emerged as a promising solution (Astolfi et al., 2020; Gupta et al., 2018; Legarreta et al., 2021). By leveraging advanced deep-learning-based architectures, these methods can efficiently learn and represent the intricate geometry of streamlines, capturing their variability while significantly reducing computational demands (Ghazi et al., 2023).

The complexity of streamlines arises from their inherently high-dimensional nature. Unlike individual points in 3D space, streamlines are sequences of points that trace complex anatomical pathways in the brain, making their representation and analysis computationally demanding, especially when large numbers are involved. To address this challenge, autoencoders (AEs), a type of unsupervised representation learning method leveraging deep learning, can be used. AEs effectively learn low-dimensional latent-space representations (embeddings) from high-dimensional streamline data. These embeddings can be directly used or fine-tuned for downstream prediction tasks.

Zhong *et al*. proposed an atlas-free method using a deep recurrent long short-term memory (LSTM)-based AE to embed varying-length streamlines into a fixed-dimensional latent space for unsupervised clustering (Zhong et al., 2022). Trained on reference bundles from the ISMRM 2015 Tractography Challenge dataset, the method significantly improved tractogram parcellation. Legarreta *et al*. introduced FINTA, a convolutional AE that filters out implausible streamlines, thereby enhancing tractography reliability (Legarreta et al., 2021). This method mapped streamlines into a latent space where similar data clustered as neighbors, enabling efficient classification based on Euclidean distances rather than computationally extensive geometric calculations. They subsequently proposed CINTA, an AE-based framework for clustering unlabeled tractography data, and demonstrated high performance on synthetic datasets and anatomically consistent clustering in human data (Legarreta et al., 2022). Building on these ideas, GESTA employed a generative AE to synthesize anatomically plausible streamlines for individual bundles while enforcing structural, connectivity, geometric, and fixel-based constraints (Legarreta et al., 2023). These developments led to FIESTA, a fully automated framework that integrates FINTA (filtering), CINTA (bundle segmentation), and GESTA (streamline generation) into a unified pipeline for robust tractography segmentation (Dumais et al., 2023). Feng *et al*. recently introduced Macrostructure-Informed Normative Tractometry (MINT), a framework based on variational AEs (VAEs) that jointly models white matter macrostructure, specifically streamline geometry, and microstructure, represented by diffusion-derived metrics. After bundle segmentation, MINT reconstructs both streamline geometry and diffusion features, quantifying deviations from normative white matter organization through reconstruction errors. These errors are then aggregated into along-tract profiles for subsequent statistical analysis (Feng et al., 2025).

While these studies demonstrate that the latent representation can preserve meaningful information about streamline geometry, bundle organization, and white matter microstructure, its potential for whole-brain mapping and visualization of white matter anomalies remains limited. Notably, no framework currently provides a voxel-wise representation of white matter deformation based on streamline-level geometric deviations. Furthermore, visualizing abnormalities at the level of individual streamlines in whole-brain tractograms is challenging due to the high density and complex spatial arrangement of white matter fibers, which results in substantial visual occlusion. Implementing such a method could facilitate intuitive visualization and characterization of tumor-induced white matter alterations. In the context of gliomas, where tumor-induced deformations disrupt the white matter, mapping these abnormalities can provide valuable insights for surgical planning by identifying distant regions affected by the tumor’s impact. Furthermore, quantifying white matter deformation may provide valuable biomarkers of structural disruption with potential relevance for clinical outcomes and prognosis. Because such a representation is derived solely from streamline coordinates, latent-space distances quantify geometric differences between reconstructed streamlines; we therefore use the term deformation to refer to streamline-geometric deviations from reference trajectories, which in glioma patients are expected to capture tumor-associated displacement, distortion, and infiltration of white matter pathways.

To address this gap, we develop a framework for detecting and mapping white matter deformations directly from whole-brain tractography. Unlike previous approaches, we first train a 1D convolutional AE trained on synthetically generated random streamlines to learn low-dimensional latent representations of streamline trajectories. This approach provides a robust, flexible representation of streamlines that is independent of brain anatomy, specific tractography algorithm, or fiber-tracking parameters. Latent representations from healthy brains are then clustered to define normative patterns of white matter organization, which serve as a reference for quantifying streamline deviations. In contrast to previous methods that depend on predefined bundles, the proposed framework derives normative white matter organization directly from whole-brain tractography, eliminating reliance on bundle segmentation. Finally, streamline-level deviations were then transformed into voxel-wise anomaly maps, enabling intuitive visualization and spatial characterization of white matter deformation. By leveraging this compact, interpretable AE representation, the proposed framework enables ranking and visualization of white matter anomalies, thereby translating structural deformations into meaningful image-based representations.

## 2. Methods

### 2.1. Overview of the pipeline

The overall pipeline for anomaly detection and mapping is illustrated in Figure 1A and consists of three main steps:

1. AE training: Train a 1D convolutional AE (Hinton & Salakhutdinov, 2006; Masci et al., 2011) to learn low-dimensional latent representations of streamlines in human brain tractograms using a synthetic dataset of 100 million (M) randomly generated streamlines.
2. Clustering reference latent representation: Project reference tractogram streamlines into the latent space using the trained encoder, and cluster the resulting embeddings to represent normative streamline patterns.
3. Deformation mapping: Project test tractogram streamlines into the latent space, calculate structural anomaly scores based on each streamline’s embedding distance to the reference clusters, and visualize these scores as an image.

**Figure 1.**
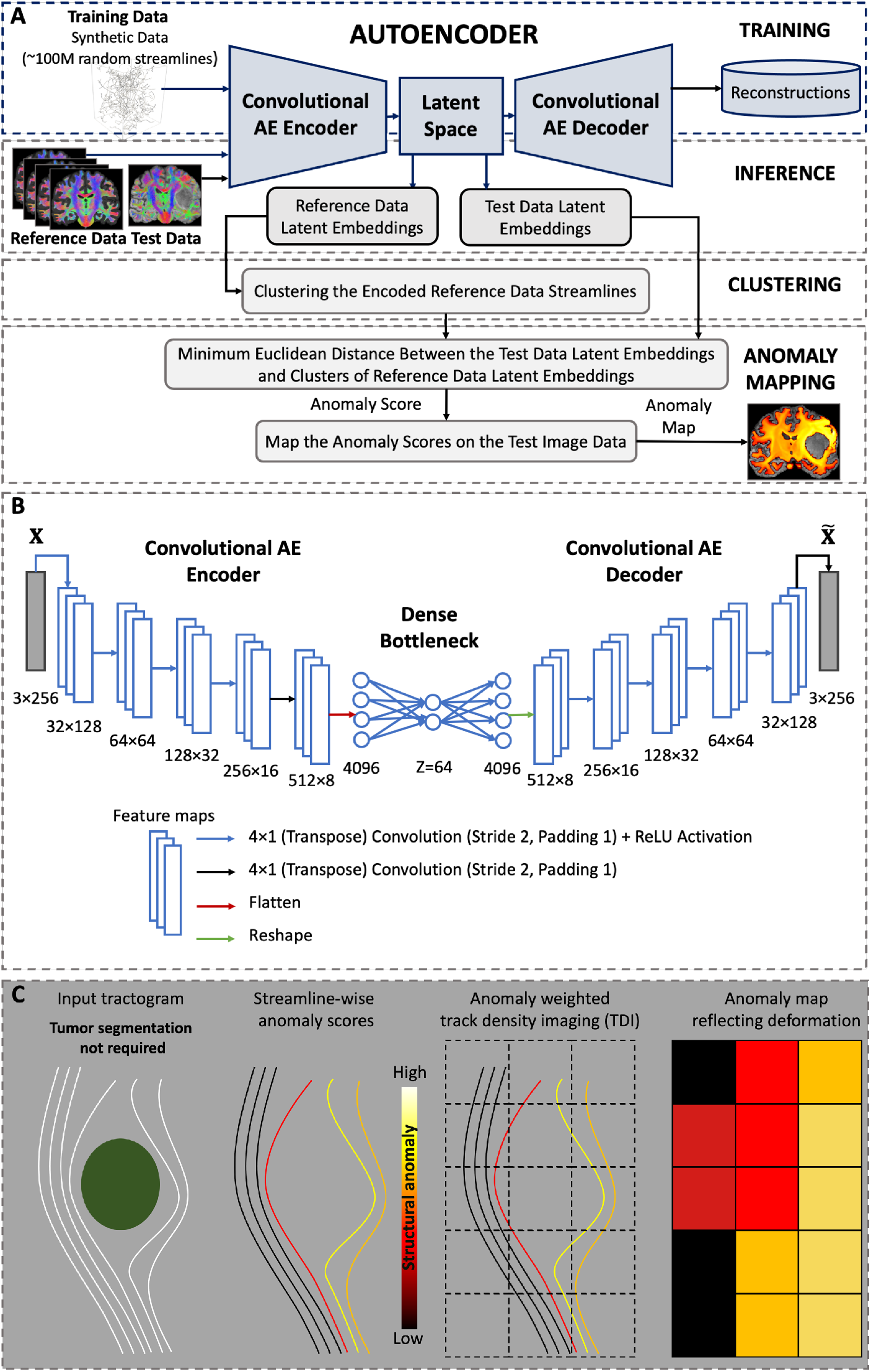
Pipeline flowchart of the proposed convolutional AE-based framework for structural anomaly detection and mapping. **(A)** The structural anomaly detection and mapping pipeline is depicted, including AE training, inference on reference and test tractograms, clustering of reference streamline embeddings, and deformation mapping. **(B)** The architecture of the 1D convolutional AE is shown. The network comprises five layers in both the encoder and decoder, with a fixed 64-dimensional dense bottleneck. Both the encoder and the decoder use a kernel size of 4, a stride of 2, and a padding of 1. ReLU activation functions are used throughout the convolutional and deconvolutional layers, except in the final layer of each module. **(C)** The schematic illustrates the deformation mapping and visualization procedure. The tumor (green) causes varying degrees of displacement and deformation in adjacent white matter pathways. Each test streamline is projected into the latent space, where its structural anomaly score is calculated as the minimum Euclidean distance to the reference cluster centers. Streamline-wise anomaly scores are aggregated into a voxel-wise image using anomaly-weighted track density imaging, with scores weighted by streamline length and normalized by local tract density. The resulting anomaly map reveals spatial patterns of structural deformation associated with the tumor. The color bar represents the degree of structural anomaly (mm), from low (black) to high (yellow).

The following sections describe synthetic streamline generation, AE training and evaluation, the reference clustering strategy, and the deformation mapping and visualization.

### 2.2. Synthesized random streamlines: Generation and methodology

We generated a synthetic dataset comprising 100M random streamlines. These streamlines were generated randomly in the MNI space, MNI152 6^th^-generation (Grabner et al., 2006), using a modified version of the Trekker software (https://dmritrekker.github.io/), which enabled the parallel transport tractography (PTT) algorithm to propagate towards a random direction at each step without using any fiber orientation from dMRI data (Aydogan & Shi, 2021). Each streamline was generated with fiber tracking parameters randomly selected from the following uniform distributions: minimum radius of curvature (mCurv) ∼ *U*(0.2,2), step size ∼ 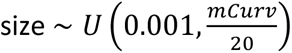, and length ∼ *U*(1,400).

### 2.3. Autoencoder architecture and training

The proposed convolutional AE model processes streamlines resampled to 256 vertices, ensuring consistency across both short and long streamlines (local cortical U-fibers vs. long-range association tracts). This choice also aligns well with previous studies that standardized streamlines for deep learning applications (Dumais et al., 2023; Legarreta et al., 2021, 2022, 2023). The AE consists of an encoder with five 1D convolutional layers and a decoder that mirrors the encoder with five deconvolutional layers. Both the encoder and decoder use a kernel size of 4, a stride of 2, and padding of 1. ReLU activation functions are applied throughout the convolutional and deconvolutional layers, except for the final layer of each module. The bottleneck is implemented as a fully connected dense layer with a fixed latent dimension of 64. For training, we used an L1 (absolute error) loss function and the Adam optimizer (Kingma & Ba, 2014). The model was trained for 10 epochs on a synthetic dataset of 100M randomly generated streamlines (80/20 split) and a batch size of 128 streamlines. The learning rate and weight decay were fine-tuned using the validation set. The final learning rate and weight decay were set to 10^-3^ and 10^-4^. All experiments were conducted in Python 3.9 (Pilgrim & Willison, 2009) using PyTorch (Paszke et al., 2019) on an NVIDIA A100 GPU (40 GB memory). Figure 1B illustrates the detailed network architecture.

### 2.4. Distance comparison in native and latent streamline spaces

We compared pairwise streamline distances in the native and low-dimensional latent streamline space to assess how well the learned embeddings preserve streamline geometry. In the native streamline space, we computed streamline distances using two measures: the Minimum Direct Flip (MDF) distance (Garyfallidis et al., 2012) and the Hausdorff distance (Huttenlocher et al., 1993). Both measures are orientation-invariant in the native streamline space. MDF achieves this by evaluating both the original and reversed point correspondences and retaining the minimum distance. In contrast, the Hausdorff distance is inherently orientation-invariant because it is defined on the set of streamline points and is independent of point ordering. To achieve the same orientation invariance in latent streamline space, we defined the streamline distance as the minimum Euclidean distance between their latent representations, considering both original (*z*) and flipped (*z*′) versions. Importantly, flipping a streamline in our model results in a corresponding flip in the latent space. Therefore, we measured the distance between two streamlines *S*_1_ and *S*_2_ as:

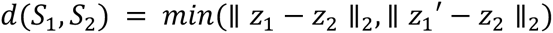

where *z*_1_ and *z*_2_ denote the latent representation of original streamlines, *S*_1_ and *S*_2_, respectively, and *z*_1_′ denotes the latent representation of flipped *S*_1_. Here, ∥⋅∥2 denotes the Euclidean norm.

For distance comparison analysis, we independently sampled 10,000 pairs of streamlines from each of four datasets, yielding 40,000 pairs in total. The datasets comprised (i) the random streamline test set, as explained earlier (Section 2.2), and (ii) Human Connectome Project (HCP) (Van Essen et al., 2012) tractograms (118 subjects) reconstructed with parallel transport tractography (PTT) (Aydogan & Shi, 2021), spherical deconvolution streamline (SD_STREAM) (J. Tournier et al., 2012), and improved fiber orientation distribution, second order (iFOD2) (J. D. Tournier et al., 2010) algorithms. For each pair, we calculated the Hausdorff and MDF distances in native streamline space and the Euclidean distance between the corresponding latent representations. Separate linear regressions were used to quantify the relationship between the native- and latent-space distances for each streamline set, with model fit summarized using the coefficient of determination (*R²*). We used a global, model-specific scaling factor of *α* to express latent-space distances in approximate physical units (mm). To assess this factor, we fitted an additional regression constrained through the origin between native-space Hausdorff distances and latent-space Euclidean distances for 10,000 pairs from the random-streamline test set. The zero-intercept constraint ensures that zero latent-space distance corresponds to zero scaled distance. We performed all analyses using Python 3.9 and the Scikit-learn library (Pedregosa et al., 2011).

### 2.5. Processing of the reference streamlines

We registered the reference streamlines to MNI space using a rigid transformation (dataset-specific details are provided in the relevant sections), resampled them to 256 vertices, and projected them into the latent space with the trained AE model. Next, we clustered the encoded representations with an in-house implementation of the leader algorithm (Hartigan, 1975; Murtagh & Contreras, 2012). This algorithm processes streamline latent representations in a single pass. For each streamline (and its flipped version), it calculates the Euclidean distance to all existing cluster leaders. The streamline is assigned to the corresponding cluster if the minimum distance is within a user-defined threshold provided in streamline space (mm), which is converted to the latent equivalent using scaling factor *α* (see Section 2.4). Otherwise, the latent representation of the streamline creates a new cluster and becomes its leader. Cluster centers (CLCs) organized in a kd-tree (Bentley, 1975) do not contain the flipped representations, since flipping is handled when computing the distance. We repeated this process until all clusters and their centers were computed, as shown in Algorithm 1. We implemented the clustering in C++ to enhance scalability (handling large datasets), improve computational efficiency, and optimize memory usage. Our implementation is based on our computational neuroimaging library, nibrary (https://github.com/nibrary/nibrary), which can sequentially read and process large numbers of streamlines stored in tractogram files within a folder. We used dynamic arrays and hash maps for memory management, and distance calculations were parallelized. Cluster management utilized the nanoflann library (Blanco & Rai, 2014; Muja & Lowe, 2009), which provides a fast algorithm for nearest-neighbor matching in high-dimensional spaces using kd-trees. Overall, C++ optimizations enabled scalable and efficient clustering of large numbers of streamlines.

#### Algorithm 1 Preprocessing and Clustering of Reference Streamlines

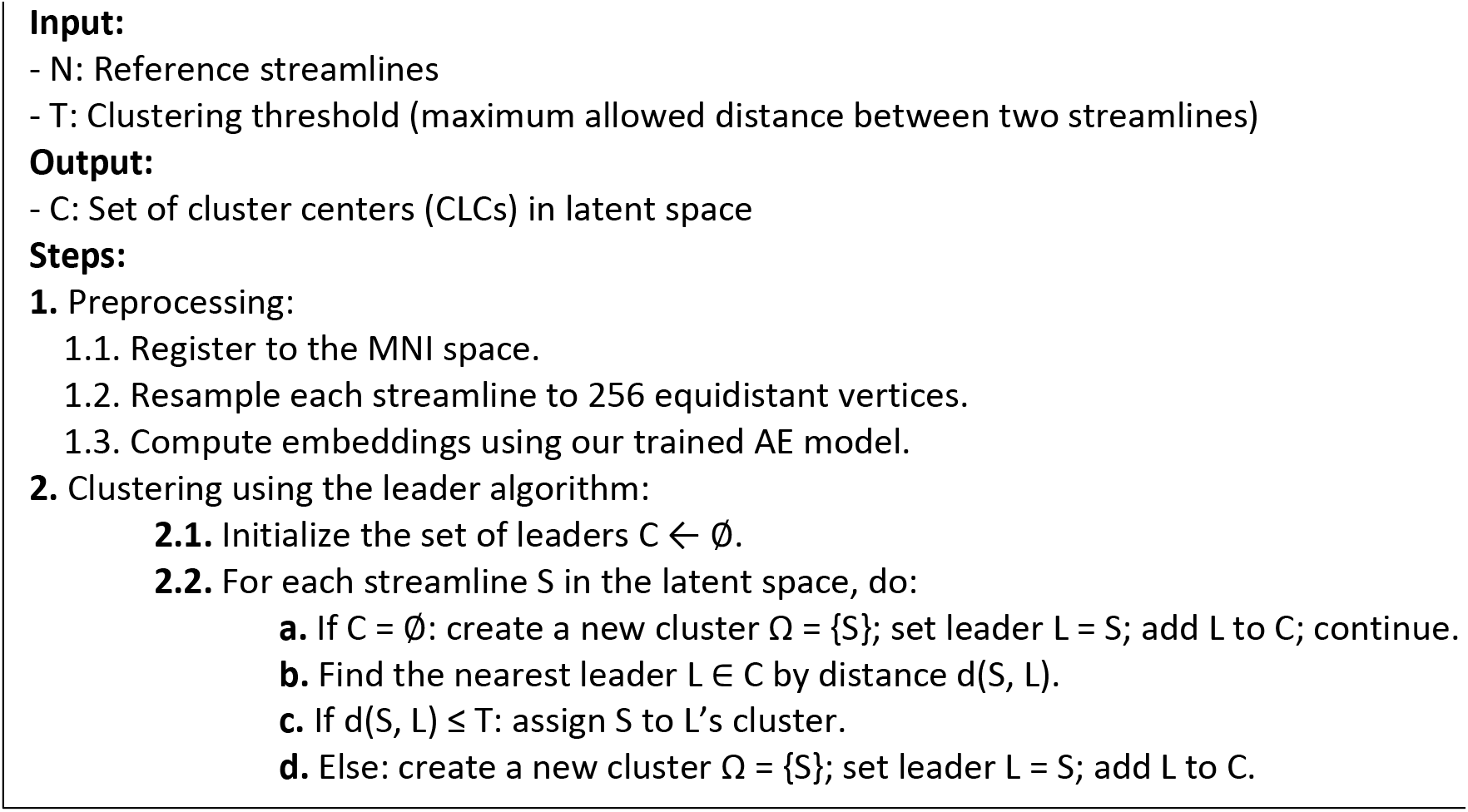

### 2.6. Deformation mapping and visualization

Using reference CLCs that represent normal streamlines, we calculated an anomaly score as a measure of deviation from normal to quantify geometric deformations for each test streamline in three steps: (i) Project the streamline into its latent space, (ii) Measure its distance to all reference CLCs, and (iii) Assign the minimum distance as the anomaly score. Formally, for a streamline *S*, with latent representation *z*, and a set of CLCs {*c*_1_, …, *c_k_*}, the anomaly score is defined as:

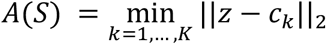

where ∥⋅∥_2_ denotes the Euclidean norm. The resulting anomaly score reflects the degree of geometric deviation from the reference population, with higher values indicating greater deformation.

To generate a voxel-wise representation of structural deformation, streamline-level anomaly scores were converted into a scalar anomaly image (Figure 1C). For each voxel *v*, anomaly scores were aggregated using a length-weighted contribution from all streamlines traversing that voxel as:

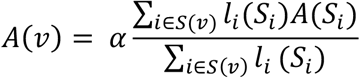

where *α* is the global, model-specific scaling factor that converts latent-space distances to physical units (mm) (see Section 2.4), *S*(*v*) denotes the set of streamlines passing through the voxel *v*, *A*(*S_i_*) is the anomaly score of streamline *S_i_*, *l_i_*(*S_i_*) *is* the total length of the segment of *S_i_* that pass through voxel *v*, i.e., the length-weighted track density image (TDI). The resulting anomaly map offers a spatial representation of structural deformation across the brain while preserving information from individual streamlines. In this framework, deformation refers specifically to geometric deviation of reconstructed streamline trajectories from reference trajectories.

### 2.7. Evaluation of the deformation mapping pipeline across multiple datasets

We evaluated the pipeline using the simulated FiberCup phantom dataset, the ISMRM 2015 tractography challenge dataset (Fillard et al., 2011; Maier-Hein et al., 2017; Neher et al., 2017), and data from 118 WU-Minn HCP subjects (Van Essen et al., 2012). We also examined clinical feasibility in two glioma datasets acquired with *high-quality* multi-shell and *clinical-quality* single-shell dMRI.

#### 2.7.1. FiberCup phantom dataset

We used the simulated FiberCup dMRI dataset (Fillard et al., 2011; Neher et al., 2017), which contains seven bundles and a ground-truth of 8,250 streamlines (https://www.nitrc.org/frs/?group_id=627). We clustered the latent representations of the ground-truth streamlines using our approach at four clustering thresholds (90 mm, 20 mm, 10 mm, and 5 mm). For each threshold, we used the ground-truth streamlines as input and generated anomaly images based on the corresponding set of cluster centers. This setup allowed us to evaluate how clustering threshold parameters influence anomaly detection. Moreover, we generated the FiberCup ground-truth streamlines length-weighted TDI.

#### 2.7.2. ISMRM 2015 tractography challenge dataset

We used the ISMRM 2015 tractography challenge dataset (Maier-Hein et al., 2017) to provide a controlled and more realistic benchmark for evaluating anomaly mapping. This dataset is synthesized from an HCP subject using 200,433 ground-truth streamlines and 25 ground-truth fiber bundles. The dMRI data have a 2 mm isotropic voxel dimension, 32 gradient directions, a b-value of 1000 s/mm^2^, and two b0 images with reverse phase encodings (More detailed information about the dataset is available at https://tractometer.org/ismrm_2015_challenge/data).

We designed three experiments. First, we used the ISMRM ground-truth streamlines as the reference and the test tractograms to generate anomaly maps, in which no anomalies were expected (similar to the FiberCup evaluations). We applied clustering thresholds of 5, 10, 20, and 50 mm to latent representations to assess their influence on anomaly mapping results. Moreover, we generated the ISMRM ground-truth streamlines length-weighted TDI.

The second experiment examined the variability caused by different fiber tracking parameters. Specifically, we examined how fiber orientation distribution (FOD) amplitude thresholds used to generate tractograms affect the resulting anomaly images. To this end, we denoised the dMRI data using the MPPCA algorithm (Veraart et al., 2016) and removed Gibbs ringing artifacts following the method described by Kellner et al. (Kellner et al., 2016). Distortion corrections were performed using the topup (Andersson et al., 2003) and eddy_openmp (Andersson & Sotiropoulos, 2016) tools from FSL (S. M. Smith et al., 2004). FODs were computed using the algorithm proposed by Tran and Shi (Tran & Shi, 2015), represented as coefficients of 8^th^-order spherical harmonics. Fiber tracking was carried out using the PTT algorithm implemented in Trekker (Aydogan & Shi, 2021). To evaluate the impact of the tracking threshold, we systematically varied the minimum FOD amplitude threshold “*minFODamp*” in Trekker from 0.01 to 0.10 across tractograms, while all other tracking parameters remained at their default values (minimum length: 40 mm; maximum length: 250 mm). Next, we quantified bundle overreach (false positives) and bundle overlap (true positives), and we generated anomaly maps for each tractogram using CLCs derived from the ground-truth latent representation with a clustering threshold of 5 mm.

In the third experiment, we used the ISMRM ground-truth corticospinal tract (CST) and applied synthetic deformations with amplitudes of 1, 2, 5, and 10 mm using a custom-designed script. For each streamline, a smooth non-rigid deformation was introduced by applying a half-sinusoidal displacement to the x-coordinate according to:

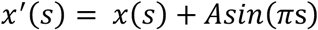

where *x*(*s*) and *x*′(*s*) denote the original and deformed x-coordinates, respectively, s∈[0,1] denotes the normalized position along the streamline and *A* represents the deformation amplitude in mm. This approach produces a smooth, non-rigid bending with zero displacement at the endpoints and maximum displacement at the streamline midpoint. We then generated anomaly maps for each deformed CST using reference CLCs derived from the undeformed ISMRM ground-truth tractogram in the first experiment, with a clustering threshold of 5 mm.

#### 2.7.3. Human Connectome Project dataset

We evaluated our pipeline on large-scale, *high-quality* dMRI data from healthy subjects in the WU-Minn HCP Data-1200 release to establish a normative benchmark for anomaly detection in healthy and disease-affected brains. We included participants who completed the full 3T imaging protocol during Q1 and Q2. dMRI data were acquired with 1.25 mm isotropic resolution using a multi-shell scheme (b-values=1000, 2000, 3000 s/mm²; 90 directions per shell; 18 b0 volumes) with both *l*eft-*r*ight and *r*ight-*l*eft phase encoding directions. We used the minimally preprocessed HCP dataset, which includes motion and eddy current correction, susceptibility distortion correction using fieldmaps, b0 intensity normalization, brain extraction, and registration to each participant’s native structural space.

We estimated tissue-specific response functions for white matter, gray matter, and cerebrospinal fluid (CSF) (Dhollander et al., 2019) using “*dwi2response*”. These response functions were then used to estimate white matter FODs using multi-shell multi-tissue constrained spherical deconvolution (CSD) with “*dwi2fod msmt_csd*” (Jeurissen et al., 2014; J.-D. Tournier et al., 2004). Whole-brain tractograms were then reconstructed for each subject using three tractography algorithms:(i) PTT (Aydogan & Shi, 2021), (ii) SD_STREAM (J. Tournier et al., 2012), and (iii) iFOD2 (J. D. Tournier et al., 2010). For SD_STREAM and iFOD2 tractography, we used MRtrix3 (J.-D. Tournier et al., 2019), while PTT was executed using Trekker. For all algorithms, we generated 10 M streamlines per subject, with minimum and maximum lengths of 2.5 mm and 250 mm, respectively, using the default FOD amplitude cutoff and step-size parameters. For iFOD2 and SD_STREAM, streamlines were dynamically seeded based on FOD amplitude and propagated using anatomically constrained tractography (ACT) (R. E. Smith et al., 2012) with five-tissue-type (5TT) masks, which were generated via the Hybrid Surface and Volume Segmentation (HSVS) framework (Fischl, 2012; Jenkinson et al., 2012; R. Smith et al., 2020; S. M. Smith et al., 2004). Backtracking was enabled, and streamlines were cropped at the gray matter–white matter interface to ensure anatomically plausible cortical terminations (R. E. Smith et al., 2012, 2015). SD_STREAM tractography used 4^th^-order Runge-Kutta integration to prevent curvature overshoots (Basser et al., 2000). For PTT, streamlines were randomly seeded across the whole white matter and propagated using ACT with the same 5TT masks as in MRtrix3 methods.

We established reference CLCs by randomly selecting 91 of 118 HCP subjects as reference tractographies, representing the “*normal*” cohort. For each tractography method, we clustered the latent representations of all streamlines (91 subjects × 10M streamlines per subject = 910 M streamlines) using a clustering threshold of 20 mm, resulting in method-specific CLCs (PTT_CLC, SD_STREAM_CLC, and iFOD2_CLC) that served as references for anomaly mapping. The remaining 27 subjects formed the test cohort. For each test subject, we randomly sampled 1M streamlines to compute anomaly scores and generate anomaly maps for each of the three reference CLC sets. This yielded nine conditions per subject, covering every combination of the three input methods and three reference CLC sets: (iFOD2, iFOD2_CLC), (iFOD2, PTT_CLC), (iFOD2, SD_STREAM_CLC); (PTT, PTT_CLC), (PTT, iFOD2_CLC), (PTT, SD_STREAM_CLC), (SD_STREAM, SD_STREAM_CLC), (SD_STREAM, iFOD2_CLC), and (SD_STREAM, PTT_CLC).

To visualize the distributions of anomaly scores, we extracted voxel-wise anomaly scores from the anomaly maps for each HCP test subject. These scores were then used to characterize the anomaly-score distributions across the nine input-reference combinations.

We assessed the effects of input and reference CLC tractography methods on subject-level mean anomaly scores using a linear mixed-effects model (LMM). The model included input tractography method, reference CLC, and their interaction as fixed effects, with subject as a random intercept to account for repeated measures. We assessed the significance of the interaction using a likelihood-ratio test comparing maximum-likelihood models with and without the interaction. Following a significant interaction, paired Wilcoxon signed-rank tests were used to compare reference CLC methods within each input tractography method and input tractography methods within each reference CLC. Holm correction was applied across all 18 post-hoc comparisons. For each comparison, we report the median paired difference with a bootstrap 95% confidence interval and the matched-pairs rank-biserial correlation (effect-size measure). Statistical significance was set at *p* < 0.05. Further details regarding model estimation, diagnostic assessment, confidence-interval estimation, and effect-size calculation are provided in the Supplementary Materials.

Moreover, for each tractography method, we implemented linear regression and compared the subject-level mean anomaly scores between pairs of reference CLCs. We reported the *R²* to show the strength of the linear association.

#### 2.7.4. Glioma patient dataset

To investigate the anomaly patterns in glioma patients, we analyzed data from two patient cohorts: one comprising two patients with *clinical-quality* single-shell dMRI data and the other with three patients with *high-quality* multi-shell dMRI data. Both cohorts included patients with left perisylvian gliomas (age range: 26–77 years), WHO grade 2-4, classified according to the 2021 WHO Classification of Tumors of the Central Nervous System (Louis et al., 2021).

### High-quality

dMRI data were acquired from patients using a Siemens 3T PRISMA scanner with a 64-channel head coil at BCAN (Charité Campus, Berlin, Germany). The multi-shell acquisition included both *a*nterior-*p*osterior (AP) and *p*osterior-*a*nterior (PA) phase-encoding directions, with 1.5 mm isotropic spatial resolution. The protocol included 14 b0 volumes and 92 diffusion-weighted directions at each of b=1500 and b=3000 s/mm², resulting in 184 diffusion-weighted directions across all b-values. Imaging parameters were as follows: echo time (TE) = 89.20 ms, repetition time (TR) = 3230 ms, matrix size = 140 × 140, number of axial slices = 92, and multiband acceleration factor of 4. We preprocessed the dMRI data using MRtrix3 (J.-D. Tournier et al., 2019) using the following steps: denoising (Veraart et al., 2016), removal of Gibbs ringing artifacts (Kellner et al., 2016), correction for subject motion (Leemans & Jones, 2009), eddy current distortions (Andersson et al., 2017), and susceptibility-induced distortions (Andersson et al., 2003) using the FMRIB Software Library (Jenkinson et al., 2012). This was followed by bias field correction with ANTs N4 (Tustison et al., 2010). Subsequently, we generated FODs as previously described for HCP data.

### Clinical-quality

dMRI data were acquired from patients using a Siemens 3T Skyra scanner with a 32-channel head coil at Charité University Hospital (Berlin, Germany). The single-shell acquisition had 2 mm isotropic resolution, with 1 b0 volume as baseline and 45 diffusion-weighted directions at b=1000 s/mm². Preprocessing followed the same steps as the *high-quality* multi-shell data. Tissue response functions for white matter, gray matter, and CSF were estimated similarly to HCP and *high-quality* patient data. Although three tissue response functions were estimated, we used only white matter and gray matter-like isotropic compartments to compute FODs with two-tissue CSD as recommended for single-shell data (Jeurissen et al., 2014; J.-D. Tournier et al., 2004). In this context, the gray matter-like isotropic compartment accounts for non-white matter signal, thereby preventing isotropic contributions from being misinterpreted as fiber orientation and improving the reliability of white matter FODs for tractography.

We reconstructed whole-brain tractograms for all patient data (both *high-quality* and *clinical-quality*) following the previously outlined steps for HCP data, with one modification. Before performing the FreeSurfer parcellation required for HSVS-based 5TT segmentation, we applied virtual brain grafting (KUL_VBG tool) to replace pathological tissue with normal-appearing tissue (Radwan et al., 2021). We then performed HSVS-based 5TT segmentation on the virtually grafted T1-weighted (T1w) or T1w-contrast-enhanced (CE) image and edited the 5TT image to incorporate the lesion mask as an additional, pathological tissue class, thereby removing tractography constraints within that region. Tractograms were registered to MNI152 space using Trekker’s “*trekker transform*” command, which employs rigid transformation through ANTs (Avants et al., 2011).

We computed anomaly scores and generated maps for all patients’ whole-brain tractographies reconstructed with PTT, SD_STREAM, and iFOD2 methods. For all analyses, we used reference CLCs derived from 91 HCP subjects with the PTT algorithm. Moreover, to quantify differences between patients and the healthy cohort, we calculated the Wasserstein-1 (W1) distance (Villani, 2009) between the empirical distributions of voxel-wise anomaly scores for each patient and the corresponding pooled HCP distribution from the 27 HCP test subjects using the same input tractography method. Each voxel was assigned equal probability mass within its respective distribution, such that each empirical distribution had a total probability mass of one. Larger W1 values indicate greater distributional dissimilarity between the patient and HCP distributions.

## 3. Results

### 3.1. AE model evaluation

We trained an AE model with a 64D latent space using a synthetic dataset of random streamlines (Methods 2.2–2.3). The distance analysis involved 10,000 randomly selected streamline pairs sampled from (i) the random streamlines test set (Figure 2A) and (ii) the HCP tractograms (118 subjects) reconstructed with three different tractography algorithms: PTT (Figure 2B), SD_STREAM (Figure 2C), and iFOD2 (Figure 2D). Latent-space Euclidean distances were strongly associated with native-space MDF and Hausdorff distances across all four datasets (*R^2^* = 0.82-0.98; Figure 2A–D, left panels: MDF vs. Euclidean distance; right panels: Hausdorff vs. Euclidean distance), which indicates that streamlines in proximity within the latent space are also close in the native streamline space. This supports the use of the embeddings to represent pairwise geometric differences. Point color indicates the combined streamline length, calculated as √(*L₁*² + *L₂*²), where *L₁* and *L₂* are the lengths of the paired streamlines, assigning lighter colors to pairs with a larger combined length magnitude. This visualization illustrates how streamline-pair length relates to pairwise distance. Moreover, the zero-intercept regression yielded a slope of 0.08 mm per latent-space unit (R² = 0.88). Throughout the analyses, we used α = 0.08 as the global scaling factor to express latent-space distances in approximate physical units (mm).

**Figure 2.**
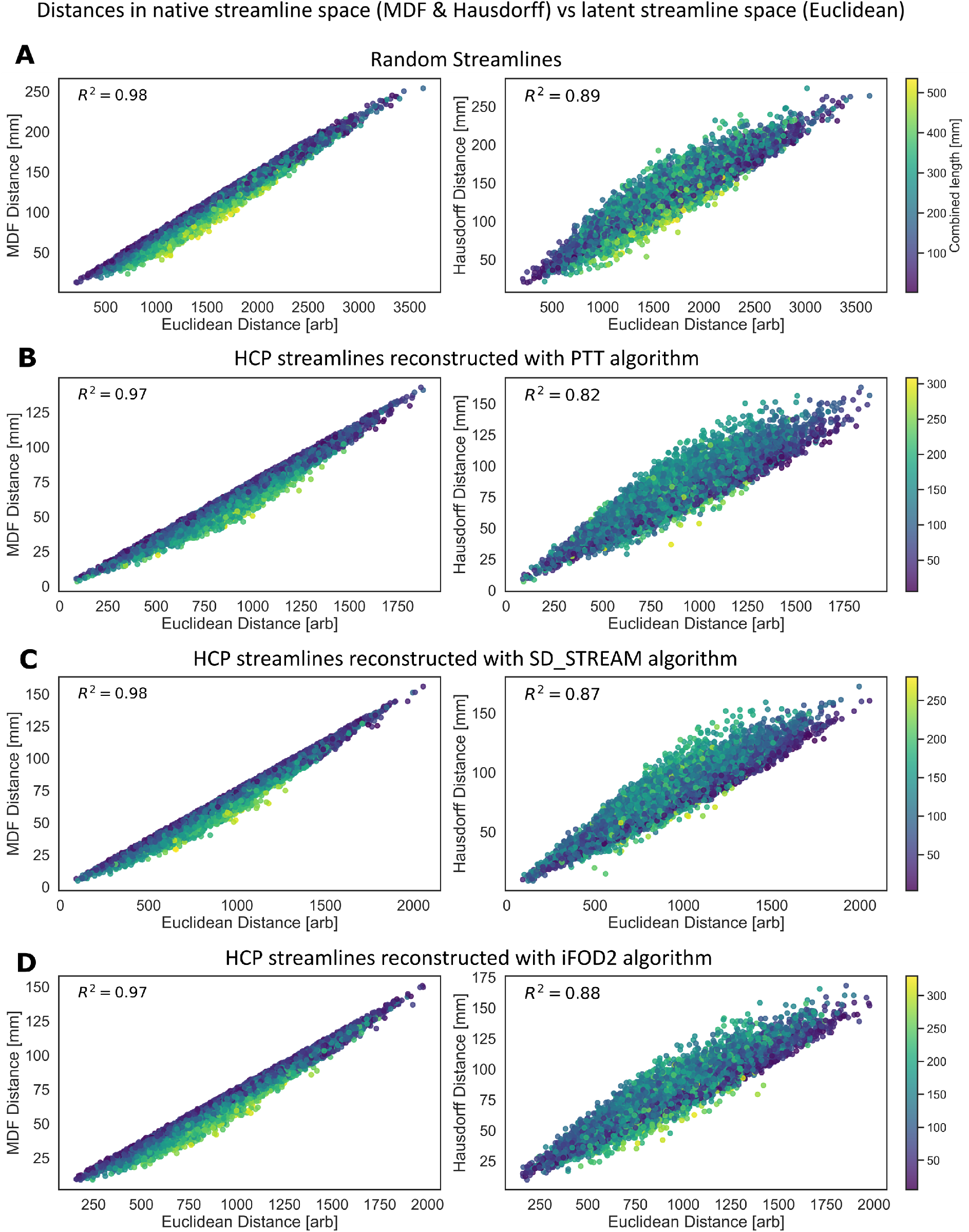
Distance comparison between the native and latent streamline spaces. The plots display the relationship between the distances in the native and 64D latent space, based on 10,000 randomly selected streamline pairs from: (A) a test set from a random streamlines dataset, the HCP dataset comprising 118 subjects with (B) PTT, (C) SD_STREAM, and (D) iFOD2 tractography methods. The x-axis shows the Euclidean distance between pairs in the latent space, while the y-axis shows the MDF distance (left plot) and Hausdorff distance (right plot) for the corresponding streamline pairs in the native streamline space. Point color encodes the combined length of the streamline pair √(L₁² + L₂²), where L₁ and L₂ are the lengths of the two streamlines. The color bar shows the corresponding metric values, with lighter colors indicating longer streamline pairs.

### 3.2. Deformation mapping and visualization: FiberCup phantom dataset

Analysis of the FiberCup ground-truth dataset showed that lowering the clustering threshold produced a more detailed anatomical representation, reflected in a greater number of representative CLCs. Reducing the threshold from 90 mm to 20 mm, 10 mm, and 5 mm increased the number of CLCs from 2 to 7, 9, and 33, respectively. The TDI served as a reference for the underlying streamline distribution and highlighted regions of high streamline density, such as fiber crossings (Figure 3). In our proposed method, structural anomaly scores are normalized by the local TDI to account for variations in streamline density and avoid preferential weighting of higher-density regions. The resulting structural anomaly maps (Figure 3) showed progressively lower anomaly scores as the clustering threshold decreased, approaching zero at the lowest thresholds. These findings suggest that the clustering threshold directly influences the method’s sensitivity, with lower thresholds enhancing sensitivity to subtle structural deviations.

**Figure 3.**
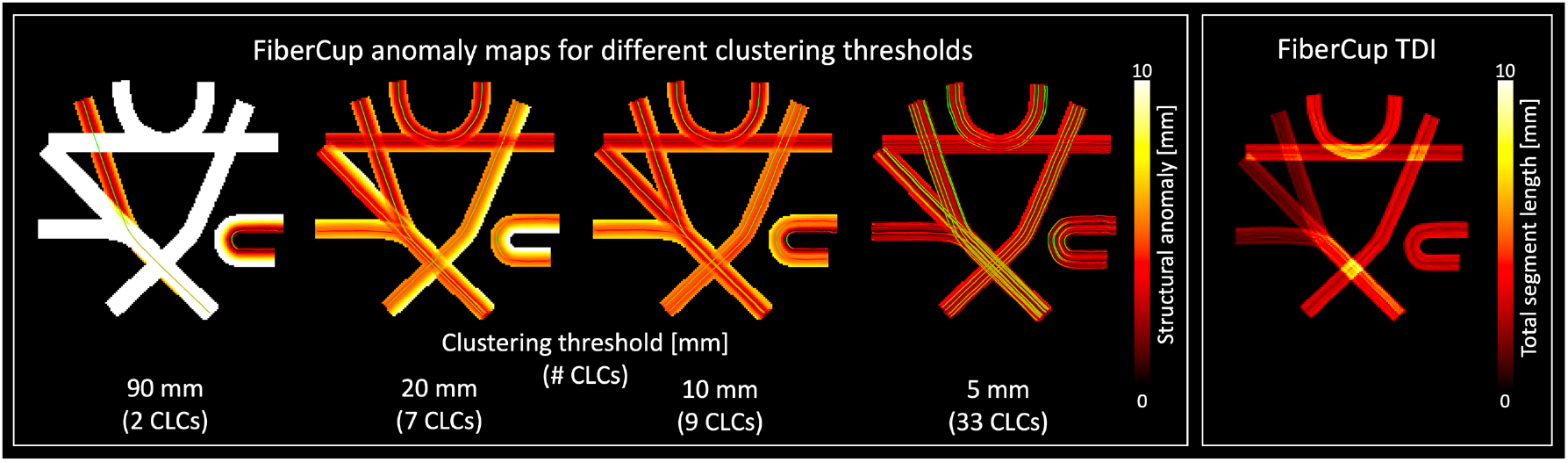
Structural anomaly analysis for FiberCup ground-truth dataset. FiberCup ground-truth streamlines were encoded using the trained AE model and subsequently clustered with thresholds of 90 mm, 20 mm, 10 mm, and 5 mm, yielding 2, 7, 9, and 33 CLCs, respectively. The left four panels display the corresponding structural anomaly maps with decoded CLCs overlaid. The anomaly color bar indicates the structural anomaly scores (mm), with higher values indicating a greater degree of anomaly. The right panel presents the FiberCup ground-truth TDI, with the color bar indicating total segment length per voxel (mm).

### 3.3. Deformation mapping and visualization: ISMRM 2015 tractography challenge dataset

Using the ISMRM 2015 tractography challenge dataset, we found that the resulting anomaly images were influenced by both the clustering threshold applied to the reference data’s latent representation and the tractography parameters used to generate the input tractograms.

In the first experiment, where no anomalies are expected, varying the clustering thresholds (5 mm, 10 mm, 20 mm, and 50 mm) resulted in 6403, 1286, 241, and 32 CLCs, respectively. Figure 4A shows the reconstructed CLCs and corresponding anomaly images, with each row representing a different threshold. The TDI maps presented alongside the anomaly maps serve as a reference for the spatial distribution and local density of reconstructed streamlines. Higher TDI values indicate regions with greater streamline density. As the clustering threshold decreased, the reconstructed CLCs yielded increasingly detailed representations of tract anatomy, and the corresponding anomaly maps exhibited lower anomaly scores. Since the anomaly scores were normalized by the local TDI, these changes cannot be attributed solely to variations in streamline density across anatomical regions.

**Figure 4.**
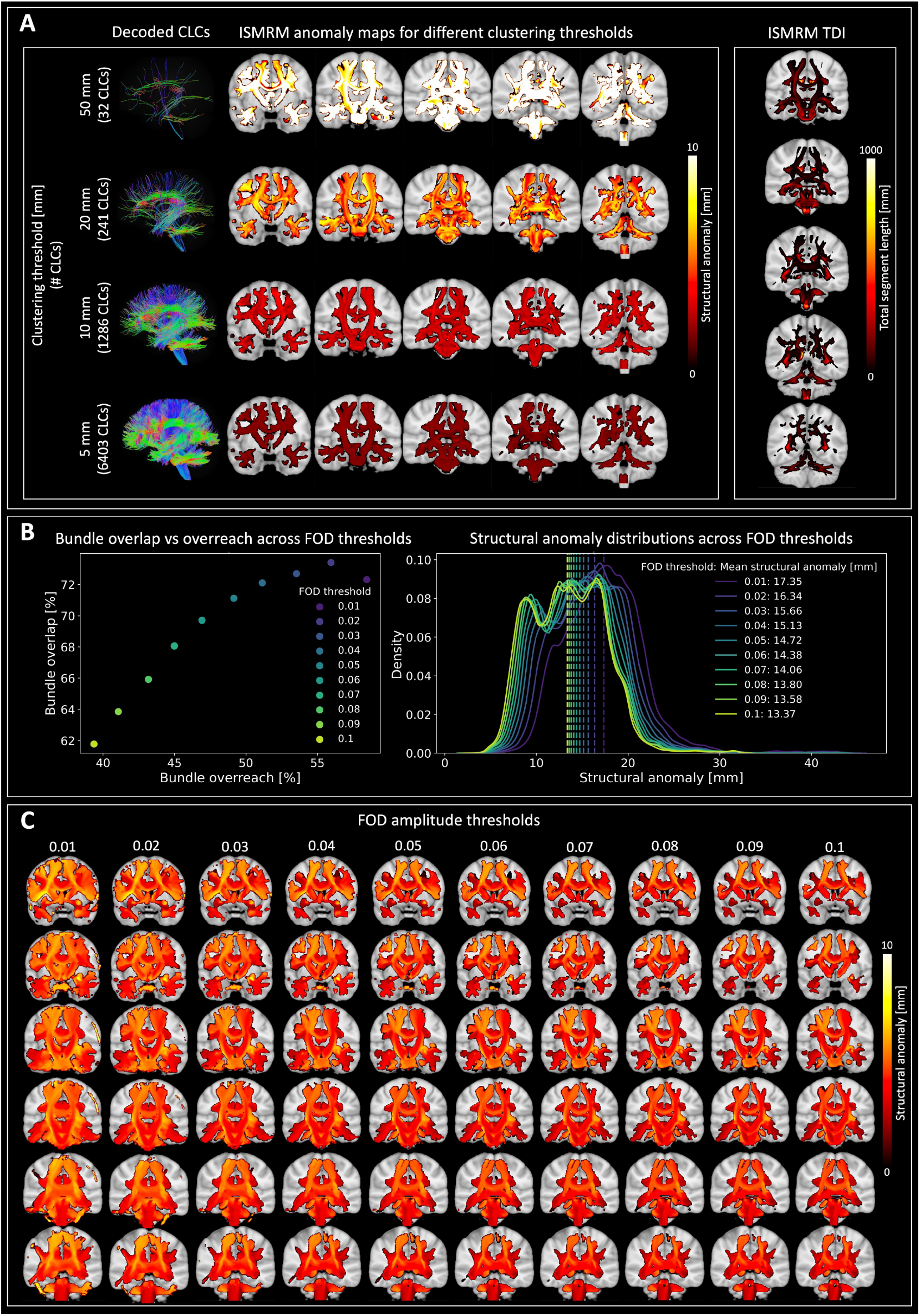
Structural anomaly analysis for the ISMRM 2015 tractography challenge dataset and tractography performance. **(A)** Decoded CLCs and corresponding structural anomaly maps derived from the ground-truth tractography using clustering thresholds of 50 mm, 20 mm, 10 mm, and 5 mm, resulting in 32, 241, 1,286, and 6,403 CLCs, respectively (from top to bottom). The rightmost column shows the corresponding TDI of the ground-truth tractography. **(B)** The left plot illustrates tractography performance across varying FOD amplitude thresholds, measured by bundle overreach versus bundle overlap. The right plot shows the PDFs of anomaly scores for these thresholds computed from 6,403 CLCs (5 mm distance threshold) derived from the ground-truth dataset. Vertical dashed lines indicate the mean structural anomaly for each threshold. **(C)** Anomaly maps across different FOD amplitude thresholds are displayed on representative multiple slices. The structural anomaly color bar reflects the intensity of structural anomaly scores (mm), with higher values indicating a greater degree of anomaly. The TDI color bar indicates total segment length per voxel (mm).

In our second experiment, as expected, a higher FOD threshold reduced bundle overreach (false positives) and bundle overlap (true positives) (Aydogan et al., 2023; Aydogan & Shi, 2021), as shown in Figure 4B (left). Therefore, we expect higher anomaly scores for streamlines generated with a lower FOD amplitude threshold, as these are more likely to represent false positives than streamlines obtained with a higher threshold. Figure 4B (right) displays the distribution of the anomaly scores as probability density functions (PDFs), obtained via kernel density estimation. Consistent with our expectation, we observed a decrease in structural anomaly values as the FOD amplitude threshold increased, with mean values ranging from 17.35 mm at an FOD amplitude threshold of 0.01 to 13.37 mm at 0.1. Figure 4C further illustrates the spatial distribution of these anomalies across multiple anatomical slices, showing that variations in the FOD amplitude threshold affected the localization and density of anomalous streamlines: lower thresholds produced a wider range of anomalous streamlines spread across the brain, while higher thresholds yielded sparser, more localized anomalies.

Figure 5 presents structural anomaly maps for the reference ISMRM ground-truth CST and its synthetically deformed counterparts with deformation amplitudes of 1, 2, 5, and 10 mm. The original CST showed minimal anomaly values, whereas increasing deformation amplitudes yielded progressively higher anomaly scores throughout the tract. Deformations of 1 mm and 2 mm resulted in relatively low anomaly values, while 5 mm and 10 mm deformations produced substantially elevated scores. Although displacement varied along each streamline, elevated anomaly scores appeared along the entire CST because each streamline was assigned a single score based on its full trajectory. Anomaly scores increased with the imposed deformation amplitude.

**Figure 5.**
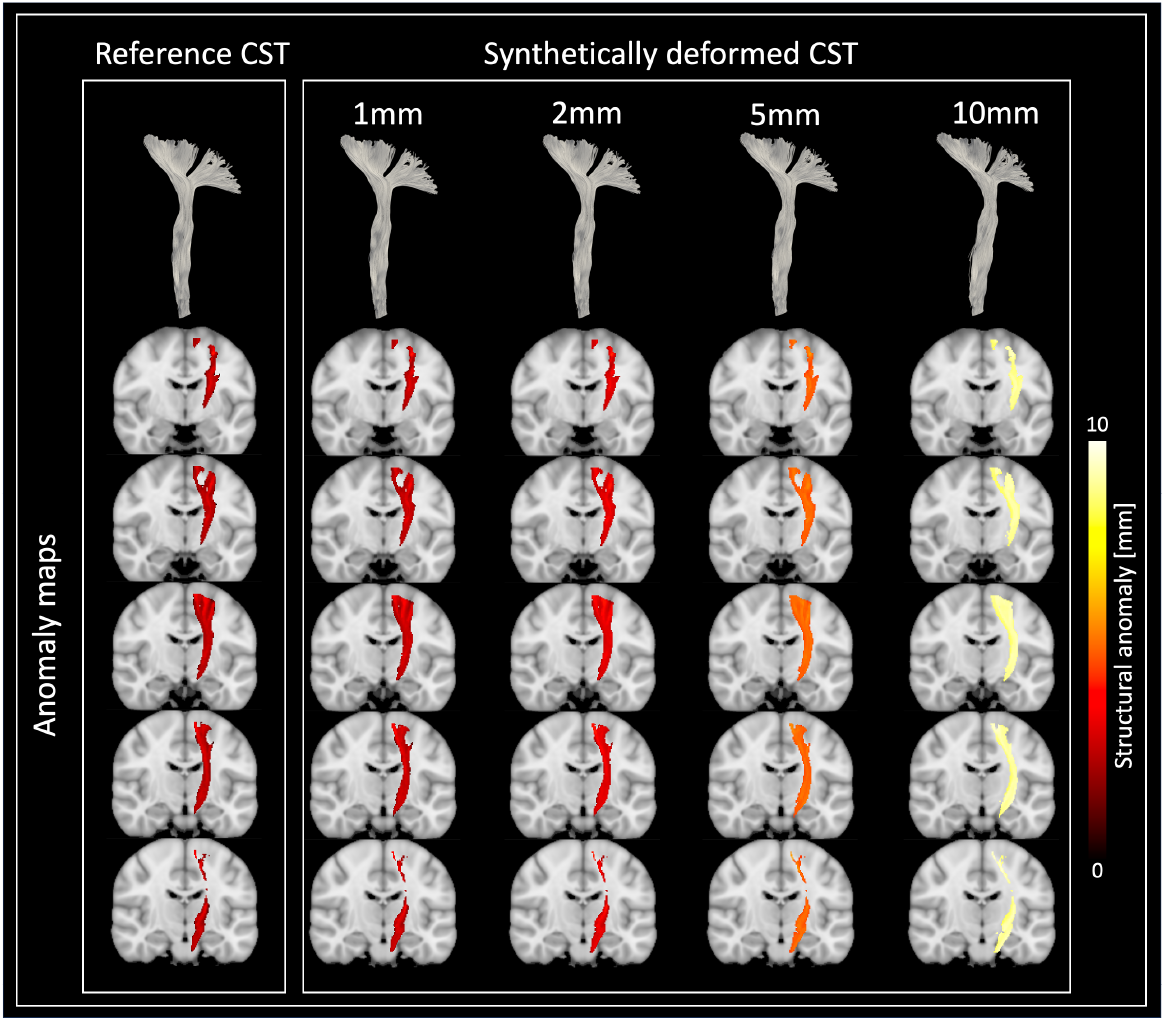
Structural anomaly maps of reference and synthetically deformed CSTs. The left panel shows the reference CST and its corresponding anomaly map. The remaining panels show anomaly maps for CSTs with synthetic deformation magnitudes of 1, 2, 5, and 10 mm. The color bar reflects the intensity of structural anomaly scores (mm), with higher values indicating a greater degree of anomaly.

### 3.4. Deformation mapping and visualization: Human Connectome Project dataset

Using the HCP dataset, we studied the robustness of our proposed approach across three tractography methods: PTT, SD_STREAM, and iFOD2. Clustering the reference cohort resulted in distinct numbers of CLCs for the three tractography methods: 495,195 for PTT, 116,451 for SD_STREAM, and 9,270,046 for iFOD2. These method-specific CLCs were subsequently used as references for anomaly mapping across the nine input-reference combinations for the test cohort.

Figure 6A shows the distributions of voxel-wise anomaly scores for the nine input-reference combinations. The first row groups the distributions by input tractography method, showing the effect of using different reference CLCs for the same input method. The second row groups the distributions by reference CLC, showing the effect of using different input tractography methods with the same reference CLC. The distributions varied across both factors, indicating that anomaly scores were sensitive to the choice of input tractography and reference CLC methods. Pooled voxel-wise mean ± SD anomaly scores were 13.15 ± 1.25, 13.76 ± 1.40, and 14.12 ± 1.30 mm for PTT input with the PTT, SD_STREAM, and iFOD2 CLCs, respectively; 12.97 ± 1.62, 12.77 ± 1.64, and 13.38 ± 1.60 mm for SD_STREAM input; and 14.79 ± 1.75, 14.74 ± 1.86, and 14.42 ± 1.43 mm for iFOD2 input, using the same reference CLC order.

**Figure 6.**
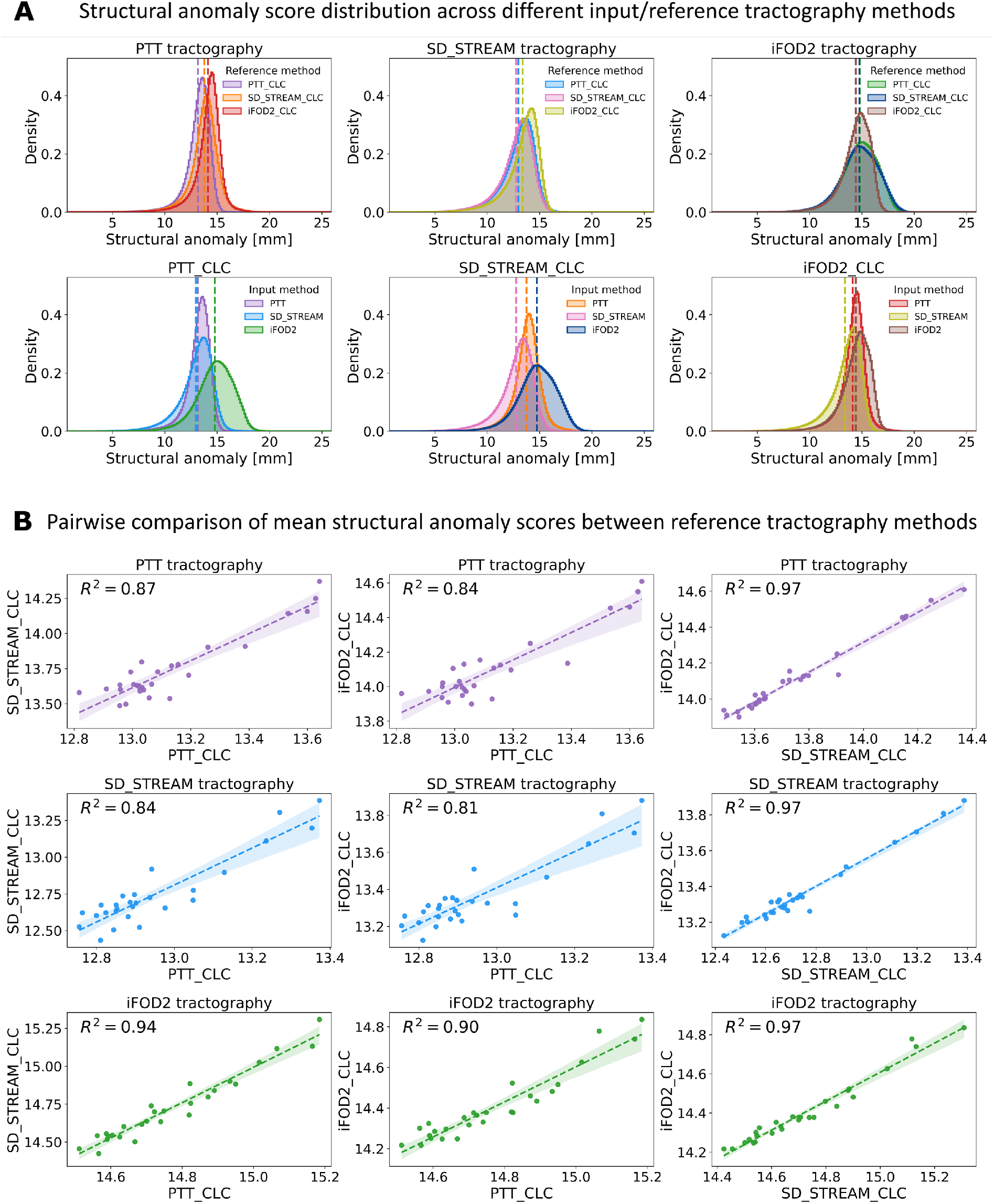
Structural anomaly analysis in the HCP test cohort. **(A)** Distribution of voxel-wise structural anomaly scores pooled across all 27 subjects for each input/reference tractography combination. (**A**, first row) Overlaid histograms are grouped by input tractography methods (iFOD2, SD_STREAM, PTT), showing the distributions obtained with different reference CLC methods for a fixed input method. (**A**, second row) Overlaid histograms grouped by reference tractography method (iFOD2_CLC, SD_STREAM_CLC, PTT_CLC), showing the distributions obtained with different input methods for a fixed reference. Dashed vertical lines indicate the voxel-wise mean anomaly scores for each distribution. Colors consistently identify the corresponding tractography methods across both rows. **(B)** Comparative analysis of subject-level mean structural anomaly scores between different reference tractography methods using the same input tractography method. Each row represents an input method (PTT, SD_STREAM, iFOD2), and each column shows a combination of reference tractography methods: (PTT_CLC, SD_STREAM_CLC), (PTT_CLC, iFOD2_CLC), and (SD_STREAM_CLC, iFOD2_CLC). Scatter plots include least-squares regression lines (dashed lines), with R² values displayed.

Descriptive statistics for the nine combinations of input tractography method and reference CLC method are provided in Supplementary Table S1. The linear mixed-effects model (LMM) identified a significant interaction between input tractography method and reference CLC method, χ^2^(4) = 591.62, *p* < 0.001, indicating that the effect of the reference CLC depended on the input tractography method. For PTT input, mean anomaly scores increased from PTT_CLC to SD_STREAM_CLC to iFOD2_CLC. For SD_STREAM input, the ordering was SD_STREAM_CLC < PTT_CLC < iFOD2_CLC, whereas for iFOD2 input it was iFOD2_CLC < SD_STREAM_CLC < PTT_CLC. Thus, no reference CLC method consistently produced the lowest anomaly scores across all input methods.

All 18 paired simple-effects comparisons remained significant following Holm correction (all adjusted *p* < 0.001). However, the magnitude of these differences varied considerably: absolute median paired differences ranged from 0.054 to 1.980 mm, and absolute rank-biserial effect sizes ranged from 0.704 to 1.000. The smallest median paired difference occurred between PTT_CLC and SD_STREAM_CLC for iFOD2 input (0.054 mm; individual unadjusted 95% bootstrap CI, 0.033–0.068 mm). Full model results and pairwise comparisons are detailed in Supplementary Tables S2, S3A, and S3B.

Figure 6B shows pairwise comparisons of subject-level mean anomaly scores between reference CLCs, with corresponding subjects matched across comparisons. Strong positive linear associations were observed across reference CLCs, with *R*² values ranging from 0.81 to 0.97. These results indicate that, although the choice of reference CLC affected absolute anomaly scores, subjects generally showed consistent relative anomaly-score patterns across reference methods.

### 3.5. Deformation mapping and visualization: Glioma patients

Figures 7 and 8 display T1w images alongside TDI and corresponding anomaly maps generated using each tractography method (PTT, SD_STREAM, and iFOD2) for both *high-quality* and *clinical-quality* data from glioma patients, respectively. Figure 7 shows T1w images, whereas Figure 8 shows T1w with contrast-enhanced (T1w-CE) images. Both figures also present voxel-wise anomaly-score distributions, descriptive statistics, and W₁ distances from the corresponding pooled HCP test-cohort distributions. All anomaly maps were computed using PTT_CLC as the reference; patient–HCP distribution comparisons used the same input tractography method.

**Figure 7.**
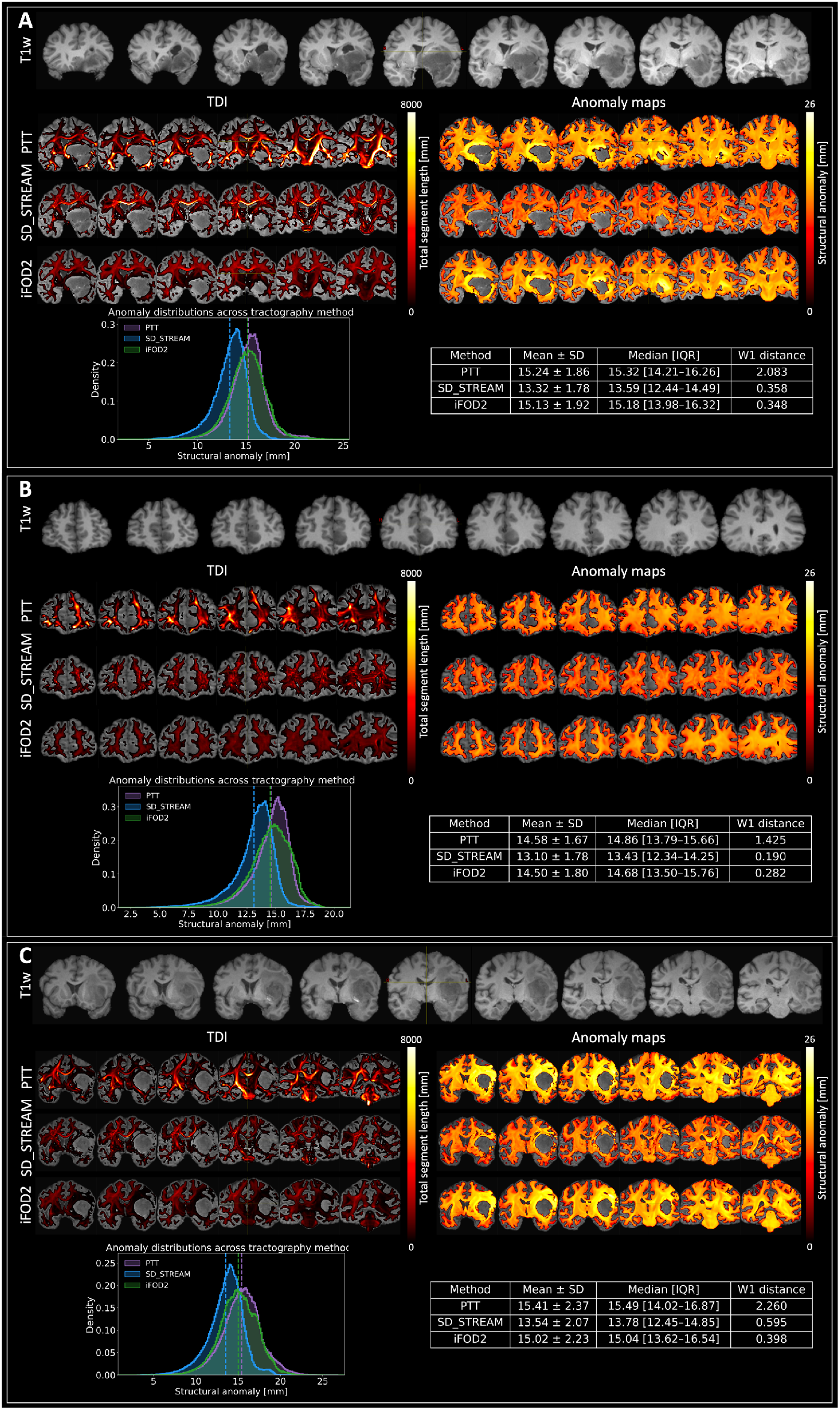
Structural anomaly analysis in left-hemispheric glioma patients with high-quality dMRI data. Results are presented for three representative cases: **(A)** a 26-year-old with diffuse astrocytoma, WHO grade 2; **(B)** a 43-year-old with diffuse astrocytoma, WHO grade 2; and **(C)** a 45-year-old with GBM. For each case, T1w anatomical images are shown in the top row. Below, TDI maps are shown on the left and the corresponding structural anomaly maps on the right, generated from tractograms reconstructed using PTT, SD_STREAM, and iFOD2 algorithms. Histograms depict the distribution of voxel-wise structural anomaly values for each tractography method, with dashed vertical lines marking the respective mean anomaly values. The accompanying tables provide descriptive statistics of the anomaly distributions and the W1 distance between each patient’s anomaly score distribution and the pooled HCP test-cohort distribution obtained using the same input tractography method. The structural anomaly color bar reflects the intensity of structural anomaly scores (mm), with higher values indicating a greater degree of anomaly. The TDI color bar indicates total segment length per voxel (mm).

**Figure 8.**
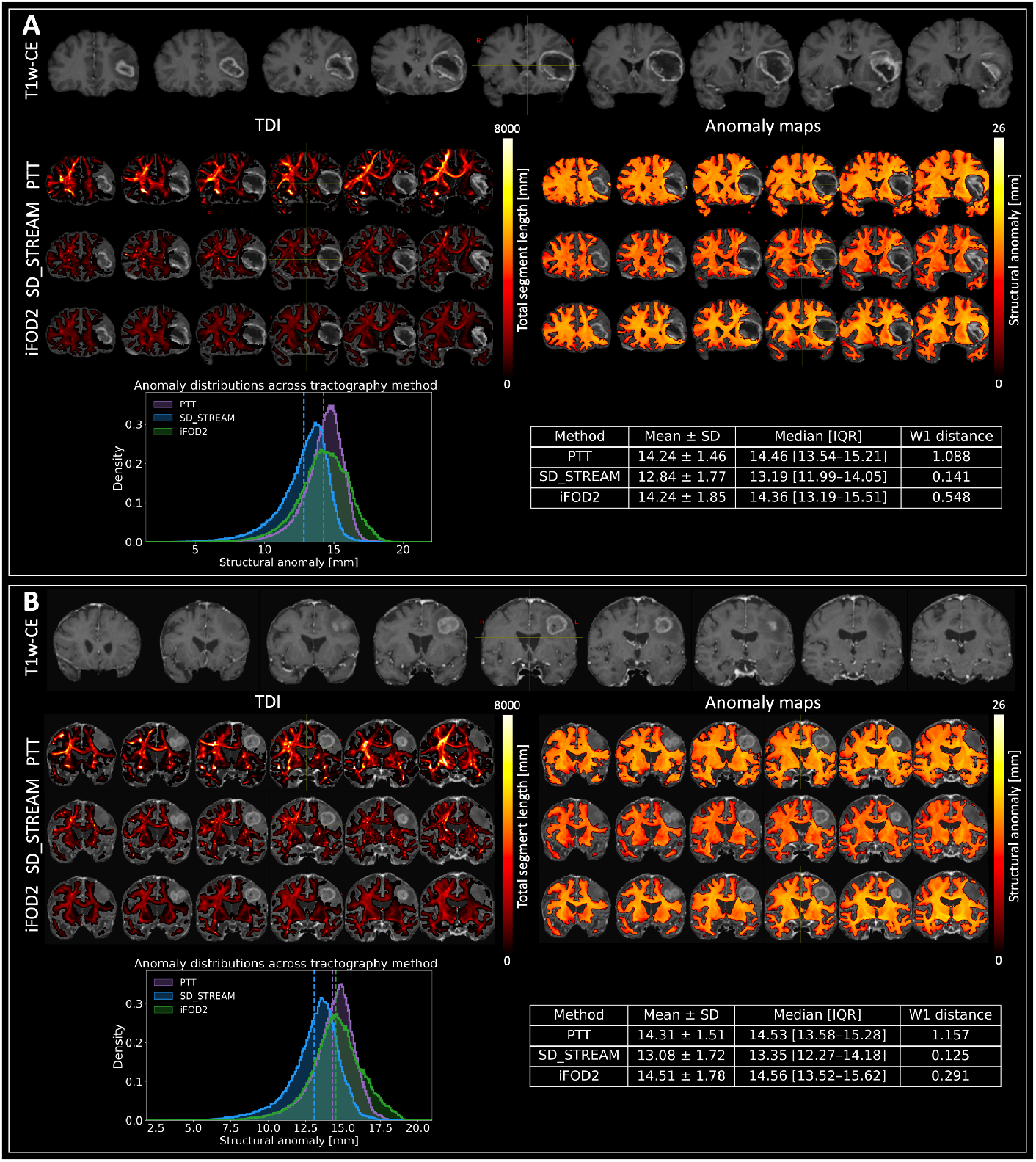
Structural anomaly analysis in glioma patients with clinical-quality dMRI data. The figure shows results for two patients: Panel **(A)** shows a 64-year-old patient with GBM, and Panel **(B)** shows a 77-year-old patient with GBM. For each case, T1w-CE anatomical images are shown in the top row. Below, TDI maps are shown on the left and the corresponding structural anomaly maps on the right, generated from tractograms reconstructed using PTT, SD_STREAM, and iFOD2 algorithms. Histograms depict the distribution of voxel-wise structural anomaly values for each tractography method, with dashed vertical lines marking the respective mean anomaly values. The accompanying tables provide descriptive statistics of the anomaly distributions and the W1 distance between each patient’s anomaly score distribution and the pooled HCP test-cohort distribution obtained using the same input tractography method. The structural anomaly color bar reflects the intensity of structural anomaly scores (mm), with higher values indicating a greater degree of anomaly. The TDI color bar indicates total segment length per voxel (mm).

Figure 7A depicts a 26-year-old patient with diffuse astrocytoma, WHO grade 2 (IDH-mutant), who presented with preoperative aphasia; Figure 7B presents a 43-year-old patient with diffuse astrocytoma, WHO grade 2 (IDH-mutant); and Figure 7C illustrates a 45-year-old patient with glioblastoma (GBM; IDH-wildtype), who also presented with preoperative aphasia. The TDI maps showed relatively lower streamline density in the peritumoral regions with clear method-dependent differences in streamline density and spatial coverage, with PTT generally providing a denser and more extensive representation than SD_STREAM and iFOD2. In contrast, the corresponding structural anomaly maps consistently identify regions of elevated anomaly values surrounding the tumor and extending into adjacent white matter. While all three tractography methods yield similar spatial patterns of structural anomalies, some differences are evident in the magnitude and spatial extent of the anomaly values.

Among the three patients and tractography methods, mean structural anomaly values ranged from 13.10 to 15.41 mm, while median values ranged from 13.43 to 15.49 mm. The distributions exhibited varying degrees of spread, with SDs between 1.67 and 2.37 mm. W1 distances from the corresponding pooled HCP distributions ranged from 0.190 to 2.260 mm. PTT consistently yielded the largest W1 distance for all three patients (2.083, 1.425, and 2.260 mm for Figures 7A, 7B, and 7C, respectively), whereas SD_STREAM and iFOD2 showed lower and variable W1 distances (0.358, 0.190, and 0.595 mm for SD_STREAM; 0.348, 0.282, and 0.398 mm for iFOD2, respectively).

Figure 8A displays a 64-year-old patient with GBM and Figure 8B a 77-year-old patient with GBM, both of whom presented with preoperative aphasia. Consistent with the findings in Figure 7, in both cases, the TDI maps showed method-dependent differences in the extent and distribution of reconstructed streamlines, with PTT generally showing more extensive coverage than SD_STREAM and iFOD2 while the structural anomaly maps showed broadly similar spatial patterns across the three tractography methods, with elevated anomaly values extending into the tumor and adjacent white matter. Differences between tractography methods were primarily apparent in the magnitude and spatial distribution of the anomaly values.

Across the two patients and three tractography methods, mean structural anomaly scores ranged from 12.84 to 14.51 mm, with median values ranging from 13.19 to 14.56 mm and SDs ranging from 1.46 to 1.85 mm. W1 distances from the corresponding pooled HCP distributions ranged from 0.125 to 1.157 mm. PTT yielded the largest distances for both patients (1.088 and 1.157 mm for Figures 8A and 8B, respectively), SD_STREAM yielded the smallest (0.141 and 0.125 mm), and iFOD2 yielded intermediate distances (0.548 and 0.291 mm).

Across both dMRI data-quality settings, PTT consistently yielded the largest W1 distance while SD_STREAM and iFOD2 showed smaller, variable distances. Similarly, PTT generally produced the highest mean and median anomaly values, whereas SD_STREAM produced the lowest, with iFOD2 generally intermediate. Despite method-dependent differences in streamline density and spatial coverage evident in the TDI maps, anomaly maps showed broadly consistent spatial distributions of elevated anomaly values across tractography methods and dMRI data-quality settings.

## 4. Discussion

In this work, we propose a novel, deep-learning-based technique that uses geometry-preserving streamline embeddings to establish a normative reference. This approach transforms latent distances from the AE into quantitative, voxel-wise anomaly images, providing a fast, computationally affordable way to study whole-brain white matter deformation across a large number of streamlines and enabling efficient computation of anomaly scores. Our method’s focus on streamline geometry offers unique insights into how pathways are displaced. This information, therefore, complements existing dMRI-based techniques such as network analysis (Bullmore & Sporns, 2009; Fekonja et al., 2022; Fornito et al., 2013; Wei et al., 2023) and along-tract microstructural profiling (Chamberland et al., 2021; Colby et al., 2012; Fekonja et al., 2021; Shams et al., 2022) A key advantage of the proposed method is that distances and anomaly scores in the latent space possess an intuitive geometric interpretation. A model-specific calibration expresses the anomaly score in approximate millimeters, although it should not be interpreted as an exact measurement of tissue displacement. This metric may have potential for broader clinical applications beyond the tumor applications demonstrated in this study. For instance, it could be applied to investigate white matter alterations in various neurological and neurodegenerative disorders, including Alzheimer’s disease, multiple sclerosis, and cerebral palsy, which are known to cause deformations in the white matter (Dou et al., 2020; Preziosa et al., 2023; Scheck et al., 2012). Each application would require disease-specific validation.

Distances in the 64-dimensional embedding space were strongly associated with native-space geometric distances (Figure 2). Using Euclidean distances between embeddings also allowed kd-tree searches for clustering and anomaly scoring. With this representation, whole-brain anomaly maps could be computed within minutes, which is difficult to achieve in the original streamline space. The biological interpretation of patient anomalies requires further validation.

The FiberCup and ISMRM experiments showed how the clustering threshold affects the reference representation and the structural details captured, with a higher number of reference CLCs revealing finer structural details. Fortunately, the intuitive nature of the physical anomaly scores (in mm) makes this parameter straightforward to tune, enabling the selection of a clinically meaningful threshold tailored for a specific application.

The HCP analysis showed that the framework could process large tractography datasets, but anomaly scores depended on both the input method and the reference CLC set. The significant interaction indicates that reference effects differed between input methods. Although subject-level scores were strongly associated across references (Figure 6B), this does not establish agreement in their absolute values. The *R^2^* values in Figure 6 indicate that SD_STREAM and iFOD2 CLCs for the HCP dataset yield anomaly scores more similar to those produced by PTT-based CLCs. Because of this difference, we used PTT_CLC for all patient examples to keep the reference fixed across analyses. The framework’s compatibility with various tractography algorithms underscores its potential as a generalizable tool for anomaly detection and scoring in the brain’s white matter. Because the reference was derived from HCP data, anomaly scores in patients may reflect not only tumor-associated deformation but also demographic, scanner, acquisition, and preprocessing differences between cohorts. Future work should evaluate matched normative references and harmonization strategies.

In glioma patients, anomaly mapping revealed higher scores near the tumor, extending into surrounding white matter and more distant white matter. These observations are consistent with previous evidence that gliomas induce white matter structural alterations detectable through dMRI (Fekonja et al., 2021, 2022; Shams et al., 2022, 2023). While the TDI maps showed relatively localized alterations in streamline density and distribution around the tumors, the anomaly maps revealed more spatially extensive deviations in reconstructed streamline geometry.

Tumor growth can cause tissue displacement and microstructural changes in surrounding white matter ((Manan et al., 2023; Pavlisa et al., 2009). Elevated values in the anomaly maps may reflect these effects, particularly in the tumor’s vicinity. In high-grade gliomas, white matter disruption may be more extensive, with structural alterations potentially extending beyond the immediate tumor region and along white matter pathways, reaching distant brain regions that were previously considered unaffected. This can show, through tractography, that gliomas can have a widespread effect on structural connectivity, affecting areas far beyond the tumor core. We interpret these patterns as potentially reflecting the combined effects of local mass effect and tumor infiltration. Mass effect may displace white matter pathways near the glioma, whereas infiltration and associated tract vulnerability may contribute to abnormalities extending distally along affected pathways. However, anomaly scores alone cannot distinguish mass effect, infiltration, edema-related tractography changes, or reconstruction errors; determining their biological basis requires independent validation. Differences among tractography algorithms likely reflect their specific biases and trade-offs, including the generation of false-positive streamlines and premature streamline termination, which may influence the apparent extent of distal abnormalities.

While connectomic analysis techniques have been used to highlight these global effects (Fekonja et al., 2022, 2025), they primarily measure changes in end-to-end connectivity. This is a significant advance over traditional dMRI-based microstructure metrics like fractional anisotropy (FA), which are sensitive only to local tissue properties (Fekonja et al., 2021; Shams et al., 2022, 2023). Our approach, on the other hand, provides a distinct and complementary perspective. By evaluating the geometric integrity of the entire streamline trajectory, our method uniquely captures how a connection’s path is altered. This offers a new layer of information beyond what is captured by either local metrics or network connectivity, deepening our understanding of the impact of glioma pathology across the whole brain. GBM cases in both data-quality cohorts showed elevated anomaly scores around the tumor and in adjacent white matter (Figures 7C and 8A–B). Within the high-quality cohort, both WHO grade 2 cases had lower mean anomaly scores than the GBM case for PTT and SD_STREAM tractography; the clinical-quality cohort contained no low-grade cases for comparison. It is important to highlight, however, that this initial work that proposes the methodological approach was not designed to test associations between anomaly magnitude and tumor grade. Therefore, the relationship between anomaly magnitude and tumor grade should be interpreted cautiously, as several factors may contribute to the observed structural deviations.

The role of edema and its impact on structural anomalies should also be taken into account. Peritumoral edema, often seen with high-grade gliomas, increases extracellular water content, which changes diffusion metrics and makes it harder to distinguish true structural changes from fluid-related effects (Farquharson et al., 2013). Consistent with this mechanism, higher anomaly scores near extensive edema should be interpreted cautiously as a mixture of geometric deformation and potential edema-induced tractography failure, rather than actual axonal disruption. Similarly, differences in tumor size, extent of infiltration, mass effect, and involvement or displacement of surrounding white matter pathways may contribute to inter-patient variability in anomaly-score distributions. Although higher anomaly values and greater distributional variability were observed in some patients with larger or higher-grade tumors, the small number of cases and the variation in tumor characteristics do not allow these factors to be considered independently. The W1 analysis provided a complementary measure of the distributional difference between each patient and the HCP test cohort. Across both high- and clinical-quality dMRI data, the method-matched PTT input/reference combination (PTT with PTT_CLC) produced the largest W1 distance in all five patients. The magnitude of the W1 distance also varied between patients, including among patients with the same histological diagnosis, indicating that the distributional deviation from the HCP cohort was not uniform across patients.

Comparison between *high-quality* multi-shell and *clinical-quality* single-shell dMRI datasets revealed differences in the sensitivity of anomaly mapping methods. *High-quality* data acquired with advanced multi-shell dMRI protocols provide greater sensitivity to subtle connectivity changes, especially in peritumoral regions where infiltration is less evident on conventional imaging. However, these regions may still have early or mild tumor infiltration. Moreover, W1 distances were generally larger in the high-quality cohort than in the clinical-quality cohort, particularly for PTT and SD_STREAM, although the relative pattern across tractography methods was preserved: PTT consistently yielded the largest W1 distances, while SD_STREAM and iFOD2 showed smaller, more variable distances, with SD_STREAM generally lowest in the clinical-quality cohort. These findings suggest that high-quality multi-shell data may capture greater distributional differences from the HCP cohort in these cases, while the consistency of the tractography-method pattern across data-quality settings supports the feasibility for the application of the anomaly mapping framework to clinical-quality single-shell data. The richer angular and diffusion-weighting information enables more accurate FOD modeling, improving detection of subtle white matter alterations (Jeurissen et al., 2014; J.-D. Tournier et al., 2019). Multi-tissue response functions for white matter, gray matter, and CSF are highly useful in tumor patients, as peritumoral regions often contain a mix of tissue types, edema, and infiltrated fibers. Multi-shell, multi-tissue CSD can separate these contributions, mitigating partial volume effects and reducing the misinterpretation of edema as tract disruption. This leads to more accurate FODs, better tractography, and more reliable structural connectivity estimations. We used a multi-tissue CSD approach with white matter for *clinical-quality* single-shell data. Although single-shell data lack the detailed angular and diffusion-weighting information of multi-shell acquisitions, including a gray matter-like isotropic compartment helps reduce partial volume effects from non-white matter tissues such as edema, CSF, or infiltrated gray matter. Nonetheless, tractography in regions affected by edema remains limited: diffusion is often less restricted in edematous tissue, causing tractography to stop farther from the tumor border. This was evident in our two GBM cases with extensive edema, where single-shell dMRI-based tractography failed to capture fibers within edematous areas. In contrast, *high-quality* multi-shell data allow more complete reconstruction of white matter pathways, even within or near edema, offering a more comprehensive view of connectivity changes in tumor and peritumoral regions. Consistent with our findings and the previous work by Farquharson et al. (Farquharson et al., 2013), multi-shell acquisition provides greater sensitivity for mapping fibers near the tumor and within edematous tissue, leading to more reliable and accurate anomaly scoring of streamlines. Nevertheless, single-shell data acquisition remains more practical and feasible in routine clinical settings.

Methodologically, although the proposed method demonstrates scalability and robustness, critical technical considerations and implications need to be addressed. A key aspect is the latent-space dimensionality. While previous studies on bundle segmentation and filtering, trained on *in vivo* human tractography, successfully used 32D embeddings using models with similar complexity as ours, our goal was to capture subtle geometric alterations caused by local streamline deformations. For our purposes, lower-dimensional embeddings were insufficient to represent these variations in randomly generated streamlines, as shown by increased reconstruction loss. We empirically found that 64D embeddings were more effective than 32D, exhibiting higher correlation with geometric distance measures (e.g., MDF and Hausdorff) in streamline space, without significantly increasing computational cost. This higher-dimensional representation thus offers an improved trade-off between expressiveness and efficiency for deformation mapping. However, we believe more complex models can achieve latent-space representations that are even more compact than 32 dimensions in the future. This would reduce the computational burden of clustering and anomaly scoring, enabling the analysis of larger cohorts with similar approaches.

Our framework computes anomaly scores by comparing a given streamline to its closest spatial neighbors in the set of reference CLCs. However, a key trade-off of our rigid registration strategy is that this spatial proximity does not guarantee true anatomical correspondence. Due to the inherent anatomical variability between subjects, the reference centers contain a wide distribution of streamline shapes. Consequently, a patient’s truly anomalous streamline may be scored as normal if it falls within this broad distribution of healthy controls, introducing noise into the anomaly calculation. While non-linear registration aims to resolve this correspondence problem, it also distorts the native streamline geometry, which is a significant limitation. Given this trade-off, we selected rigid registration as the preferred approach due to its simplicity and straightforward interpretability.

We also evaluated the effect of the clustering threshold used to define the reference CLCs. Lowering the clustering thresholds resulted in finer reference representations and increased sensitivity to subtle deviations in streamline geometry. However, excessively small thresholds produced substantially more clusters, which increased computational complexity. We selected 20 mm for the HCP reference as a practical compromise between reference detail and computational cost, and used those CLCs for the HCP and patient analyses. The optimal threshold for detecting clinically relevant changes remains to be established.

While our study investigated the variability of anomaly scores across different tractography methods, it is important to note that, for all methods, some streamlines are likely to be implausible, and many connections are inevitably missed (Maffei et al., 2022; Schilling et al., 2019). Some detected anomalies may therefore reflect reconstruction errors. Conversely, the method cannot map abnormalities in regions where no streamlines are reconstructed.

One simplification we made in our experiments was omitting quantitative tractography methods. While our computed anomaly maps are a form of track-weighted imaging, they are currently weighted solely by the geometric anomaly score. For truly quantitative results that better reflect the underlying diffusion signal, the input tractograms should be processed using techniques that filter or weight streamlines e.g., SIFT (R. E. Smith et al., 2013), SIFT2 (R. E. Smith et al., 2015), COMMIT (Daducci et al., 2015), LIFE (Pestilli et al., 2014). Our framework is fully compatible with such quantitative inputs; the final anomaly scores can be adjusted by incorporating these streamlines’ weights. Although we did not explore this integration (as our primary goal was to demonstrate the core geometric approach), it represents an important direction for future research.

Using whole-brain tractograms, we demonstrated that our approach can identify spatially distributed structural anomaly patterns in glioma patients, while the observed variability in anomaly-score distributions highlights the potential influence of tumor characteristics, tractography methods, and dMRI data quality. While the present work focused on structural alterations, future studies could investigate how these maps relate to functional outcomes and explore their potential value in preoperative assessment. Rapid, non-invasive anomaly maps may support preoperative assessment and cohort stratification once validated against clinical outcomes. Future research should test this framework in larger and more diverse patient populations, evaluate its clinical utility, and examine how structural anomaly mapping may complement other imaging modalities to guide treatment strategies.

## 5. Conclusion

This study introduces a deep learning framework for quantifying structural deformations in white matter using whole-brain tractography. The framework encodes streamline geometry into a latent representation and compares individual streamlines with a normative reference, generating voxel-wise anomaly maps that quantify structural deviations in physically interpretable units. By evaluating the entire streamline trajectory, this method offers a complementary perspective to both local diffusion metrics and end-to-end connectomic analyses, capturing alterations in white matter pathway geometry that neither approach alone may detect.

The results provide initial evidence that the proposed framework can visualize tumor-associated white matter anomaly patterns in a small set of glioma patients with varying tumor characteristics, tractography methods, and dMRI data-quality settings. While further validation in larger, more diverse cohorts is necessary, the approach’s computational efficiency and interpretability suggest its promise for studying structural alterations across neurological and neurodegenerative disorders.

## 6. CRediT authorship contribution statement

**Boshra Shams**: Data curation, Formal analysis, Investigation, Methodology, Software, Validation, Visualization, Writing – original draft, Writing – review and editing. **Lucius S. Fekonja:** Data curation, Formal analysis, Funding acquisition, Investigation, Methodology, Supervision, Validation, Visualization, Writing – original draft, Writing – review and editing. **Peter Vajkoczy:** Funding acquisition, Project administration, Resources, Writing – review and editing. **Thomas Picht:** Funding acquisition, Investigation, Project administration, Resources, Supervision, Writing – review and editing. **Dogu Baran Aydogan:** Conceptualization, Data curation, Formal analysis, Funding acquisition, Investigation, Methodology, Project administration, Resources, Software, Supervision, Validation, Visualization, Writing – original draft, Writing – review and editing.

## Acknowledgements and Funding

Thomas Picht, Lucius Fekonja, and Boshra Shams acknowledge support from the Cluster of Excellence Matters of Activity. Image Space Material, funded by the Deutsche Forschungsgemeinschaft (DFG, German Research Foundation) under Germany’s Excellence Strategy—EXC 2025-390648296. Additionally, Thomas Picht and Lucius Fekonja acknowledge support from the DFG, Computational Connectomics, SPP 2041, project number 455227709. Dogu Baran Aydogan received funding from the Research Council of Finland (grant numbers 348631 and 353798) and the Technology Industries of Finland Centennial Foundation (grant number 4302).

## Ethical approval

The study received approval from the Ethics Committee of Charité – Universitätsmedizin Berlin (EA1/016/19, EA1/129/20, and EA4/147/20) and was conducted in accordance with the Declaration of Helsinki. Written informed consent was obtained from all patients for participation and for the use of their data in research. Where applicable, separate consent for onward sharing of patient data was obtained, as detailed in the Data Availability Statement.

## Data and code availability

Patient MRI data are governed by the General Data Protection Regulation (Regulation (EU) 2016/679; GDPR), the German Federal Data Protection Act (Bundesdatenschutzgesetz; BDSG), and relevant ethics approvals and consent conditions. Consent for retrospective clinical-quality MRI data did not include public data sharing; therefore, these data cannot be made available. For the high-quality patient MRI dataset, participants provided informed consent allowing controlled data sharing within the European Union. Accordingly, pseudonymised data may be provided to qualified researchers upon reasonable request, subject to institutional approval, adherence to consent conditions, and completion of a data-use agreement. The remaining datasets are publicly available and are cited in the manuscript. The synthetic dataset is available from the corresponding author upon reasonable request. Preprocessing, processing, and analysis code are openly accessible at https://github.com/BoshraS/Anomaly_Detection.

