## Supplementary Table S1, Supplementary Table S2, Supplementary Table S3A, Supplementary Table S3B for "Autoencoder-based anomaly mapping and visualization of white matter deformation with application to glioma cases"

### Description of the linear mixed-effects model (LMM) analysis

For each of the 27 HCP test subjects, we computed the mean anomaly score for all nine combinations of input tractography method (PTT, SD\_STREAM, and iFOD2) and reference CLC set (PTT\_CLC, SD\_STREAM\_CLC, and iFOD2\_CLC). This yielded 243 observations, with one observation per subject for each condition. The subject was the unit of inference. Supplementary Table S1 summarizes the subject-level mean anomaly scores for each input–reference combination.

**Supplementary Table S1.** Subject-level mean anomaly scores for the nine input–reference combinations.

| Input | Reference | N | Mean | SD | Median | Q1 | Q3 |
| --- | --- | --- | --- | --- | --- | --- | --- |
| PTT | PTT_CLC | 27 | 13.134 | 0.227 | 13.039 | 12.999 | 13.175 |
| PTT | SD_STREAM_CLC | 27 | 13.743 | 0.235 | 13.637 | 13.601 | 13.788 |
| PTT | iFOD2_CLC | 27 | 14.102 | 0.198 | 14.022 | 13.972 | 14.132 |
| SD_STREAM | PTT_CLC | 27 | 12.952 | 0.175 | 12.893 | 12.847 | 13.012 |
| SD_STREAM | SD_STREAM_CLC | 27 | 12.750 | 0.240 | 12.674 | 12.622 | 12.760 |
| SD_STREAM | iFOD2_CLC | 27 | 13.362 | 0.189 | 13.312 | 13.255 | 13.353 |
| iFOD2 | PTT_CLC | 27 | 14.770 | 0.187 | 14.722 | 14.615 | 14.882 |
| iFOD2 | SD_STREAM_CLC | 27 | 14.725 | 0.225 | 14.678 | 14.544 | 14.861 |
| iFOD2 | iFOD2_CLC | 27 | 14.405 | 0.170 | 14.364 | 14.291 | 14.471 |

Values summarize subject-level means across 27 subjects; SD is the between-subject standard deviation; Q1 and Q3 denote the first and third quartiles. Scores are expressed in mm.

We fitted an LMM with input tractography method, reference CLC method, and their interaction as fixed effects, and a random intercept for subject to account for repeated observations. We tested the interaction with a likelihood-ratio test comparing maximum-likelihood (ML) models with and without the interaction. The test statistic was evaluated against a chi-square distribution with four degrees of freedom. Subsequently, we estimated model coefficients and variance components using restricted maximum likelihood. Model assumptions were assessed visually using a residual Q–Q plot and a plot of residuals against fitted values.

Following a significant interaction, we performed paired Wilcoxon signed-rank tests to compare reference CLC methods within each input tractography method and input tractography methods within each reference CLC. We applied Holm correction across all 18 post-hoc comparisons. Paired differences were calculated as the first method minus the second. For each comparison, we reported the median paired difference, its percentile-bootstrap 95% confidence interval (CI) based on 10,000 paired resamples, and the matched-pairs rank-biserial correlation. Statistical significance was set at $p < 0.05$.

Because voxel-wise group inference would require reliable anatomical correspondence across subjects, which may be affected by inter-individual anatomical variability and registration uncertainty, we performed the statistical inference on subject-level mean anomaly scores. The analysis therefore focused on differences in overall anomaly-score magnitude across conditions rather than spatial localization of these effects.

### Detailed LMM results

All primary random-intercept models converged. The interaction between input tractography method and reference CLC method was significant ($\chi^2(4) = 591.62$, $p < 0.001$), indicating that the effect of the reference CLC depended on the input tractography method. The estimated subject random-intercept variance was 0.037, the residual variance was 0.005, and the intraclass correlation coefficient (ICC) was 0.88. Supplementary Table S2 summarizes the interaction test, variance components, and ICC.

**Supplementary Table S2.** Primary LMM summary.

| Analysis | Statistic | df | p | Estimate |
| --- | --- | --- | --- | --- |
| **Input x Reference interaction** | $\chi^2 = 591.62$ | 4 | <0.001 | - |
| **Subject random-intercept variance** | - | - | - | 0.037 |
| **Residual variance** | - | - | - | 0.005 |
| **ICC** | - | - | - | 0.88 |

The interaction test uses ML fits; variance components and the ICC were obtained from the full model fitted by restricted ML. Variances are expressed in mm<sup>2</sup>; ICC is dimensionless.

All 18 paired comparisons remained significant after Holm correction (all adjusted $p < 0.001$). Absolute median paired differences ranged from 0.054 to 1.980 mm, and absolute matched-pairs rank-biserial correlations ranged from 0.704 to 1.000. Supplementary Tables S3A and S3B report the comparisons between reference CLCs within each input method and between input methods within each reference CLC, respectively.

**Supplementary Table S3A.** Paired comparisons of reference CLC methods within each input tractography method.

| Fixed input | Comparison | Median difference | 95% CI low | 95% CI high | Rank-biserial r | Wilcoxon W | Holm-adjusted p |
| --- | --- | --- | --- | --- | --- | --- | --- |
| **PTT** | PTT_CLC - SD_STREAM_CLC | -0.616 | -0.642 | -0.564 | -1.000 | 0 | <0.001 |
| **PTT** | PTT_CLC - iFOD2_CLC | -0.969 | -0.997 | -0.932 | -1.000 | 0 | <0.001 |
| **PTT** | SD_STREAM_CLC - iFOD2_CLC | -0.365 | -0.378 | -0.347 | -1.000 | 0 | <0.001 |
| **SD_STREAM** | PTT_CLC - SD_STREAM_CLC | 0.205 | 0.155 | 0.229 | 0.979 | 4 | <0.001 |
| **SD_STREAM** | PTT_CLC - iFOD2_CLC | -0.411 | -0.461 | -0.359 | -1.000 | 0 | <0.001 |
| **SD_STREAM** | SD_STREAM_CLC - iFOD2_CLC | -0.618 | -0.645 | -0.595 | -1.000 | 0 | <0.001 |
| **iFOD2** | PTT_CLC -<br>SD_STREAM_CLC | 0.054 | 0.033 | 0.068 | 0.704 | 56 | <0.001 |
| **iFOD2** | PTT_CLC -<br>iFOD2_CLC | 0.358 | 0.336 | 0.409 | 1.000 | 0 | <0.001 |
| **iFOD2** | SD_STREAM_CLC -<br>iFOD2_CLC | 0.321 | 0.286 | 0.362 | 1.000 | 0 | <0.001 |

Differences were calculated as the first method minus the second method (mm). CIs are individual, unadjusted 95% percentile-bootstrap intervals. Holm correction was applied jointly across all 18 comparisons in Tables S3A and S3B; all comparisons include 27 paired subjects.

**Supplementary Table S3B.** Paired comparisons of input tractography methods within each reference CLC method.

| Fixed reference | Comparison | Median difference | 95% CI low | 95% CI high | Rank-biserial r | Wilcoxon W | Holm-adjusted p |
| --- | --- | --- | --- | --- | --- | --- | --- |
| **PTT_CLC** | PTT -<br>SD_STREAM | 0.180 | 0.145 | 0.232 | 0.989 | 2 | <0.001 |
| **PTT_CLC** | PTT - iFOD2 | -1.621 | -1.688 | -1.556 | -1.000 | 0 | <0.001 |
| **PTT_CLC** | SD_STREAM -<br>iFOD2 | -1.812 | -1.838 | -1.780 | -1.000 | 0 | <0.001 |
| **SD_STREAM_CLC** | PTT -<br>SD_STREAM | 0.978 | 0.946 | 1.014 | 1.000 | 0 | <0.001 |
| **SD_STREAM_CLC** | PTT - iFOD2 | -0.975 | -1.048 | -0.917 | -1.000 | 0 | <0.001 |
| **SD_STREAM_CLC** | SD_STREAM -<br>iFOD2 | -1.980 | -2.004 | -1.923 | -1.000 | 0 | <0.001 |
| **iFOD2_CLC** | PTT -<br>SD_STREAM | 0.733 | 0.707 | 0.784 | 1.000 | 0 | <0.001 |
| **iFOD2_CLC** | PTT - iFOD2 | -0.293 | -0.368 | -0.255 | -1.000 | 0 | <0.001 |
| **iFOD2_CLC** | SD_STREAM -<br>iFOD2 | -1.040 | -1.057 | -1.012 | -1.000 | 0 | <0.001 |

Differences were calculated as the first method minus the second method (mm). CIs are individual, unadjusted 95% percentile-bootstrap intervals. Holm correction was applied jointly across all 18 comparisons in Tables S3A and S3B; all comparisons include 27 paired subjects.
